# Polycomb Repressive Complexes 1 and 2 are recruited independently to pericentromeric heterochromatin in response to hypomethylation in mouse embryonic stem cells

**DOI:** 10.1101/2025.11.14.688451

**Authors:** Silviya Dimova-Vasileva, Olga Stepanova, David Hay, Katherine E. Pickup, Philippe Gautier, Laura C. Murphy, Yatendra Kumar, Ian R. Adams, Sari Pennings, Richard R. Meehan

**Author notes:** Joint senior authors (Sari Pennings, Richard R. Meehan).

## Abstract

Pericentromeric heterochromatin (PCH) is delineated by the enrichment of repressive epigenetic modifications, specifically trimethylated histone H3 at lysine 9 (H3K9me3) and DNA methylation (5-methylcytosine), which establish and maintain a condensed, transcriptionally silenced chromatin state. Depletion of either H3K9me3 or DNA methylation in mouse embryonic stem cells (mESCs) induces a permissive chromatin configuration that permits de novo recruitment and deposition of normally excluded Polycomb Repressive Complexes 1 and 2 (PRC1 and PRC2), characterized by H2AK119ub1 and H3K27me3 modifications, respectively, at PCH. Here, we demonstrate that H2AK119ub1 and H3K27me3 are independently recruited to hypomethylated PCH using a doxycycline-inducible mESC model allowing modulation of Dnmt1 expression levels and catalytic activity. We further investigate the roles of proposed mediators of PRC1/2 targeting, including SCML2, BEND3, KDM2b, and TET enzymes, in this context, our findings indicate that neither PRC1 nor PRC2 recruitment at hypomethylated PCH depends on these factors. Additionally, our data suggest that the permissive chromatin environment resulting from DNA hypomethylation is the principal facilitator of Polycomb complex spreading, offering novel insights into the mechanisms governing epigenetic modifier dynamics and interactions during periods of DNA methylation reprogramming.

## Introduction

The pioneering cytological methods of Heitz identified the regions of post-mitotic chromosomes that stained more densely than others in interphase cell nuclei, thus defining heterochromatin and euchromatin, respectively (1, 2). These have provided cytogenetic landmarks for multiple studies in other species to identify similar structures to align the genomic landscapes (3). Now almost a century later, we have an in-depth molecular description of heterochromatic regions but still lack structural understanding of how heterochromatin is assembled and organised (4).

Early genetic experiments demonstrated that heterochromatic regions are associated with transcriptional inactivity, to an extent that genes that are relocated in its proximity via chromosomal rearrangements can exhibit variable silencing or position effect variegation (5, 6). Heterochromatin can be classified into facultative and constitutive heterochromatin (fHC and cHC). The former has the capacity to transition into a transcriptionally permissive state in specific developmental contexts including cellular differentiation (7). By contrast, cHC is a stable cytological feature across cell types, including embryonic stem cells (ESCs), which predominantly localises at (peri)centromeric regions and telomeres containing tandemly repetitive DNA sequences (8). In murine cells, pericentromeric heterochromatin (PCH) comprises arrays of satellite DNA sequences, primarily characterized by AT-richness, which facilitated their separation from the bulk genome during density gradient centrifugation (9, 10). In situ hybridisation experiments showed that these sequences map to PCH in interphase and mitotic cells, corresponding to DAPI (4ʹ, 6-diamidino-2-phenylindole) dense regions (11). Major satellite repeats are situated within pericentromeric domains, while minor satellite sequences are localised at kinetochore/centromeric regions of chromosomes (11). Perturbation of heterochromatin integrity is associated with genomic instability, demonstrating its functional importance (12).

The dense organisation of heterochromatin as suggested by DAPI staining is further substantiated by lower nuclease enzyme sensitivity of its densely positioned nucleosome arrays, and its unique biophysical properties which are thought to be the basis for transcriptional repression at or adjacent to these regions (13). This is also supported by the occurrence of DNA and histone signatures associated with transcriptional repression, notably histone hypoacetylation, enrichment for histone methylation of Suv39h1/2 enzyme dependent-H3K9me3, Suv4-20h1/2 dependent-H4K20me3, and high levels of DNA methylation at CpG dinucleotides in repeated satellite arrays (14–18).

Interestingly however, the DAPI dense appearance of heterochromatin is maintained in mESCs that lack either DNA methylation or Suv39h1/2 mediated H3K9me3 (19–21). Their absence results in the deposition of H3K27me3 at pericentromeric heterochromatin and suggests that facultative heterochromatic marks can compensate for the global loss of constitutive marks (5mC and H3K9me3). Biochemical analysis confirms that hypomethylated cHC is deficient in H3K9me3 and enriched in H3K27me3 (22). Detection of Polycomb group proteins (PcGs) at the PCH has also been observed in mESC models in context of either DNA hypomethylation or loss of H3K9me3 (21, 23). DNA methylation has been shown to prevent H3K27me3 deposition elsewhere in the genome and constrict PcGs to their target sites especially at CpG islands (CGIs) (21, 23). In its absence, H3K27me3 can be relatively enriched at new regions on a kilobase scale (24). However, how PcGs are recruited to these new regions is currently not known.

A number of additional factors enriched at hypomethylated PCH were identified that were proposed to be part of a mechanism that delivers the Polycomb Repressive Complex (PRC) 2 to PCH, which mediates trimethylation of H3K27 via the EZH2 enzyme. For instance, depletion of the transcriptional repressor *Bend3* by shRNA in the Suv39h1/2 double knockout resulted in diminished H3K27me3 immunofluorescence signal at cHC, suggesting BEND3 may be required to enable a fHC-like state at centromeric regions and contributes to its DAPI-dense appearance in mESCs (22).

Our aims for this study were, firstly, to develop a dynamic model system which can rapidly test whether a protein is responsible for driving PcG recruitment to PCH and is able to quantify the relocalisation. Secondly, we aimed to determine the timescale within which the PcGs redistribute to PCH upon loss of DNA methylation. And thirdly, we sought to evaluate the model proposing that Polycomb accessory proteins drive the relocalisation of the PRC1/PRC2 complexes upon hypomethylation.

To achieve this, we used rapid inhibition through either transcriptional repression of the maintenance DNA methyltransferase Dnmt1 or by inhibition of DNMT1 to deplete DNA methylation globally and at the PCH satellites and investigated the localisation of PRC marks using immunofluorescence and a published live-cell reporter construct for H3K27me3 (24). Using these systems, we demonstrate that H2AK119ub1 and H3K27me3 modifications are established but the marks are no longer retained at PCH when DNA methylation is restored to previously hypomethylated PHC. Using an unbiased method for the quantification of the PcG signal at the heterochromatic chromocenters, we show that PRC relocalisation occurs while the region becomes hypomethylated. We demonstrate that PRC1 and PRC2 dependent modifications are redirected to the chromocenters independently from each other and concomitantly with hypomethylation. We determine in our system that neither SCML2, BEND3, KDM2b or TET enzymes influence the redistribution of either H3K27me3, H2AK119ub1 or both, which suggests these proposed candidates are not the main drivers of PRC1/PRC2 redistribution to hypomethylated PCH. These results endorse the view that DNA methylation itself has a strong role in determining the protein composition of the PCH in mESCs and functions to reduce access to potential passenger proteins. At the same time, our results strongly support a model in which DNA methylation acts to prevent the spreading of H2AK119ub1 and H3K27me3 to cHC regions. It also suggests that it is an intrinsic property of satellite chromatin, with its positioned nucleosomes, to organize into a dense refractory chromatin fibre (25).

## Methods

### Cell lines, culture and treatments

Unless otherwise indicated, mouse embryonic stems cells were grown on 0.2% gelatin (Sigma, # G1890) coated T25 or T75 flasks at 37°C and 5% CO_2_, in Dulbecco’s Modified Eagle Medium (DMEM, Gibco, # 41965039), 15% foetal calf serum (Hyclone), 1mM non-essential amino acids (Sigma, # M7145), 1mM sodium pyruvate (Sigma, #S8636), 2mM L-glutamine (Gibco, #25030-024), 1% penicillin/streptomycin, 0.1mM beta-mercaptoethanol (Gibco, # 31350-010). WT J1, *Dnmt1/3A/3b* triple knockout (TKO) cells (26), *Ring1b*-AID (27) and *Kdm2b*^fl/fl^ cells (28) were cultured in 1000 U/ml ESGRO Leukaemia Inhibitor Factor (Millipore, #ESG1106), while 1500U/ml LIF was used for *Dnmt1*^tet/tet^ cells and their derivatives. The *Dnmt1*^tet/tet^ cell line was a gift from R. Chaillet lab (29), modified to lack neo and puro resistance cassettes. WT and *TET1/2/3* triple knockout cells (30) were cultured in DMEM with 20% foetal calf serum, 0.1mM non-essential amino acids, 1mM sodium pyruvate, 2mM L-glutamine, 0.1mM beta-mercaptoethanol, 1000 U/ml ESGRO LIF with 3µM CHIR-99021 (Stemgent, # 04-0004) and 1µM PD0325901 (Stemgent, # 04-0006).

*Dnmt1*^tet/tet^ cell line and its derivatives were treated with 2µg/ml of doxycycline to inhibit *Dnmt1* expression. Inhibition of DNMT1 protein in all other cell lines was performed using 2µM of the *Dnmt1* inhibitor (MedChemExpress, #GSK-3484862) for 4 days unless otherwise stated. *Ring1*-AID cells were treated with indole-3-acetic acid (sodium salt) (Auxin) (Cayman Chemical, #16954) to deplete RING1B protein levels for 4 days. *Kdm2b*^fl/fl^ cells were treated with 800nM (Z)-4-hydroxytamoxifen (OHT) (Sigma, #H7904) for 4 days to induce cre-mediated recombination of the CxxC domain of the *Kdm2b* gene. All cell lines were passaged ever 2-3 days using Trypsin/EDTA (Sigma, T4174).

### Generation of Knockout lines

To generate separate *Ezh2*, *Bend3* and *Scml2* knockout clones in the *Dnmt1*^tet/tet^ cell line, CRISPR/Cas9 was used to excise as much of the genes as possible in a macro-deletion. Double sgRNAs (Table 1) were cloned into pSpCas9(BB)-2A-GFP (PX458) and pSpCas9(BB)-2A-Puro (PX459) V2.0 vectors and co-transfected using FuGENE HD (Promega). Vectors PX458 and PX459 V2.0 were gifts from Feng Zhang (31). Transfected cells were selected by adding 1.8 µg/mL puromycin to the culture media for 24h. Surviving cells were plated, and individual colonies were picked and screened for the macrodeletion by PCR using primers in Table 2. Clones with homozygous deletions were expanded and used throughout the study.

**Table 1.**
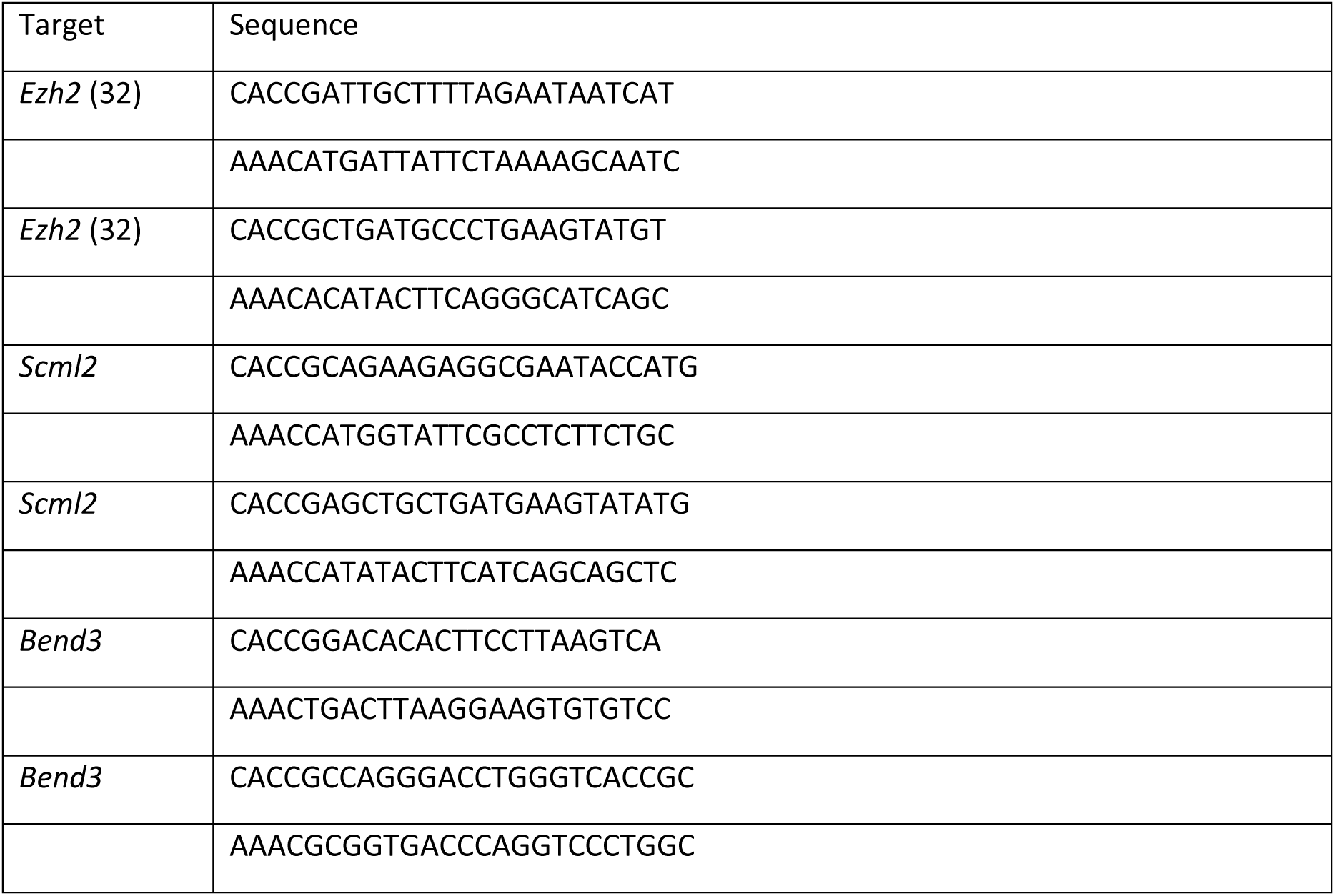
CRISPR guides sequences.

**Table 2.**
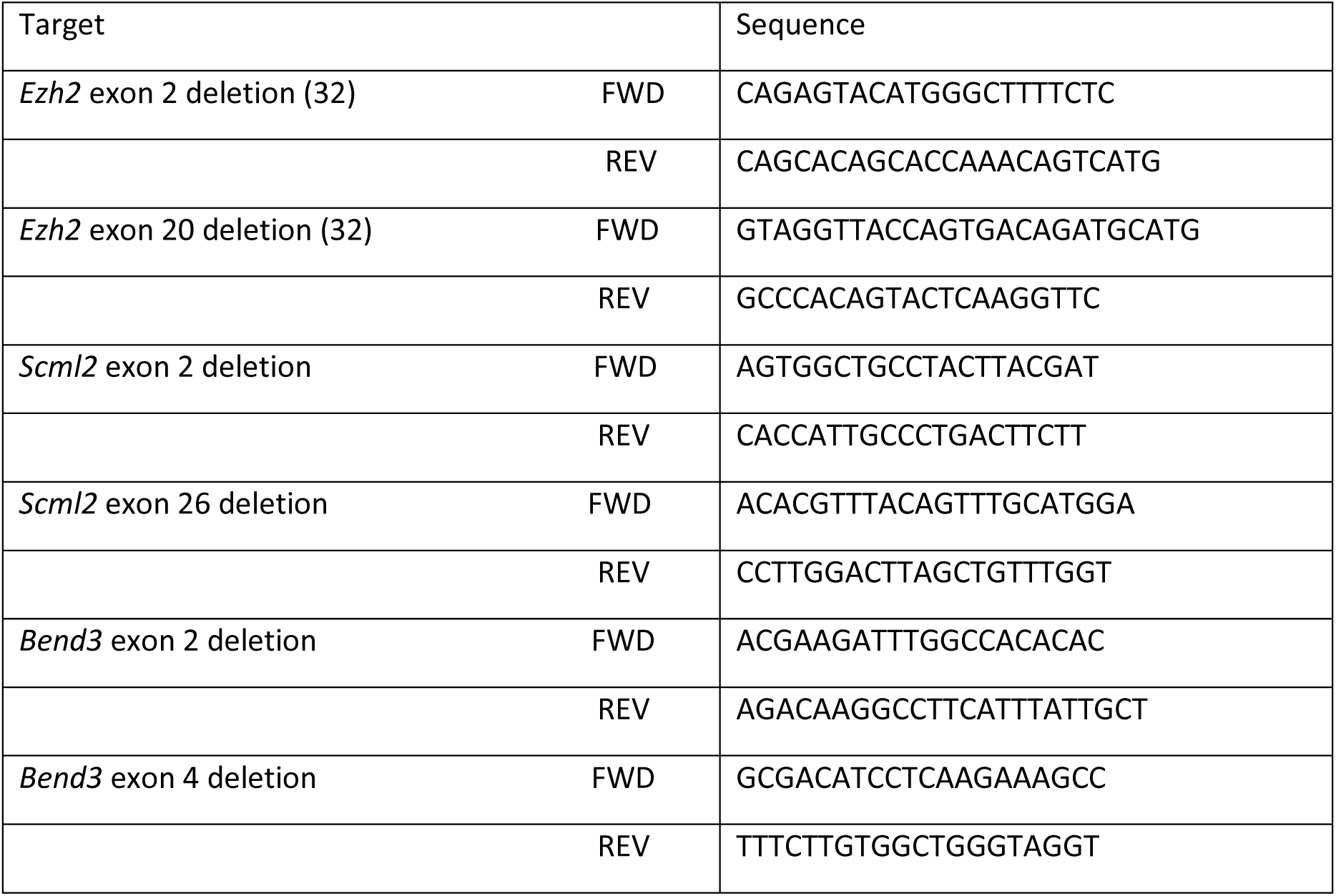
Sequencing primers.

### Gateway cloning

Target sequence from donated plasmids was amplified using primers (Table 3) that add attB overhangs in a standard PCR reaction 1X Phusion® High-Fidelity PCR Master Mix, HF Buffer (NEB, M0531S), 10mM forward and reverse primers, 1ng original plasmid (24), MilliQ water for total reaction volume −25µl. Cycling conditions were as follows: Initial denaturation (98°C – 30 sec), 30 cycles (98°C – 10 sec; 67.5°C – 20 sec; 72°C – 30 sec), Final extension (75°C – 1 min), 20°C – hold.

**Table 3.**
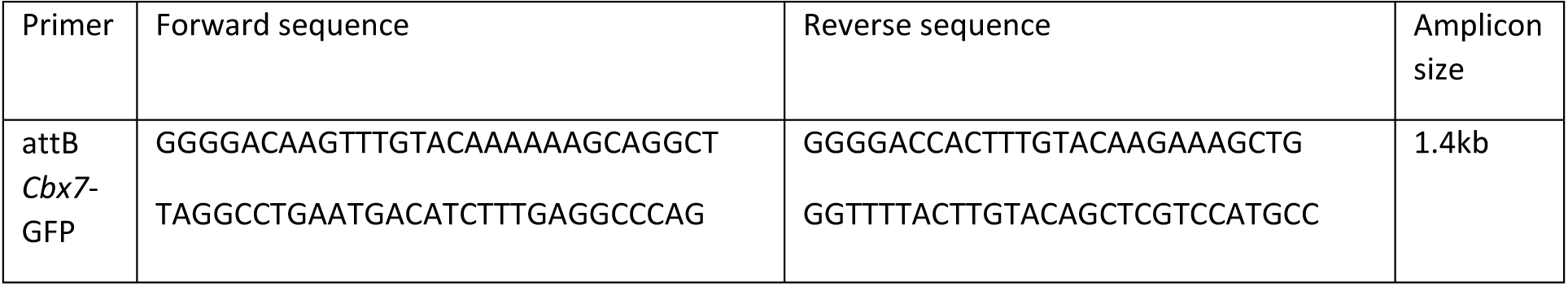
Sequences for Gateway cloning.

PCR reactions were resolved on 1% 1X TBE gel with 1x SybrSafe at 70V for 45-60 min. Bands were visualised on a Safe Imager Transilluminator (Invitrogen). The required DNA fragments were excised, gel extraction was performed using QIAquick Gel Extraction kit (Qiagen, 28704).

BP Clonase reaction containing attB containing linked DNA fragment (67.7ng), attP DonR vector (ThermoFisher, 12536017) (195ng), 1X Gateway™ BP Clonase™ II Enzyme mix (ThermoFisher, 11789020), TE buffer pH 8.0 was set up to produce an Entry clone containing the DNA fragment and left for 4h at RT after which it was inactivated by adding 2µl proteinase K and incubated at 37°C for 10 min. DH5α competent cells were transformed with 1µl of BP reaction, grown on Kanamycin plates. Single colonies were expanded in L-broth solution with Kanamycin and miniprepped using NucleoSpin Plasmid kit (Macherey-Nagel, 740588.50) following manufacturer’s protocol. Plasmids sequenced in-house. Entry clones with the correct sequence were used in LR reaction Entry clone (87.5ng), pPB-PGK-destination (150ng) (Plasmid 60436, Addgene), 1X Gateway® LR Clonase® II enzyme mix (ThermoFisher, 11791020), TE buffer pH 8.0. Reaction was performed for 1h at RT after which it was inactivated by adding 2µl proteinase K and incubated at 37°C for 10 min. DH5α competent cells were transformed with 1µl of LR reaction, grown on Ampicillin plates. Single colonies were expanded in L-broth solution with Ampicillin and plasmids extracted as above. Plasmids were sequenced in-house.

### Transfection of mESCs with PiggyBac plasmids

Cells were cultured for at least a week before transfection and were tested for mycoplasma on the day before transfection. Stably expressing cell lines were generated using a reverse transfection reaction with Fugene HD (Promega, E2311), with a ratio of Fugene HD to DNA - 4:1. The quantity of DNA in each transfection reaction was 2.25µg at ratio of 2:1 piggyBac vector containing the gene of interest to pBase transposase vector. Cells were counted at 200 000 cells and plated on a gelatinised 6 well plate and transfected in normal media containing Pen/Strep and were washed twice with PBS the following morning after which fresh media was added. The presence of fluorescent signal in appropriate wells was checked on a fluorescent microscope. Cells were cultured until fluorescent signal in the ‘No transposase’ well was lost (around 7-10 days) to ensure that all non-integrated plasmids had been expelled from the cells. The cells were then sorted for positive GFP signal using a Fluorescence-Activated Cell Sorter (FACS) as single cells into a 96-well plate and as a mixed population of cells which were then plated on a gelatinised 6 well plate. Single clones were then expanded to be used in experiments.

### Immunocytochemistry

Cells grown on 0.2% gelatin-coated glass coverslips were washed with PBS, fixed with 4% PFA for 10 min and washed 3×10 min with PBS. The coverslips were then blocked with 10% donkey serum (Sigma, #SLBT751) in 0.1% TritonX-100 for 1h, then incubated with primary antibody diluted in 1% donkey serum overnight at 4°C (Table 4), followed by 3×10 min wash with PBS. Appropriate Alexafluor-conjugated secondary antibodies (Table 5) diluted at 1:200 in 1% donkey serum were added for 1h at RT and were then washed 3×10 min with PBS. Nuclei were counterstained using DAPI (4ʹ, 6-diamidino-2-phenylindole, Invitrogen) for 10 min and washed 2× 10 min with PBS and once with H_2_O. Coverslips were then mounted on glass slides using Vectashield mounting medium (Vector labs, #H1000), sealed with nail polish and cured o/n in the dark at 4°C. Imaging was performed on a spinning disc confocal microscope (Dragonfly, Andor) using either a 40x dry or 60x oil objective, as stated in the figure legend or using an epifluorescent microscope - Zeiss AxioImager 2 (specified in figure legend). Exposure times were matched between conditions within experiments to compare signal levels. For all fixed staining images taken with confocal microscope of cells containing the Cbx7-GFP reporter, a despeckle filter (median filter) in ImageJ was applied (33). Image analysis was performed using ImageJ macros, available in GitLab repository https://git.ecdf.ed.ac.uk/meehanlab_public/heterochromatinpaper/chromocentersanalysis Data from macros was exported to excel spreadsheets. Relevant information was transferred to GraphPrism9 for data visualisation.

**Table 4.** Primary antibodies used for immunocytochemistry and western blots.

| Protein | Company | Species | Cat number | Dilution for ICC | Dilution for WB |
| --- | --- | --- | --- | --- | --- |
| DNMT1 (K18), polyclonal | Santa Cruz Biotech | goat | sc-10221 | 1:1000 | 1:1000 |
| H3K27me3, monoclonal | Cell Signalling Technologies | rabbit | C36B11 | 1:1600 | 1:1000 |
| H2AK119ub1, monoclonal | Cell Signalling Technologies | rabbit | D27C4 | 1:500 | 1:2000 |
| H3K9me3, monoclonal | Diagenode | mouse | C15200146 | 1:500 | 1:1000 |
| EZH2, monoclonal | Cell Signalling Technologies | mouse | AC22 | 1:100 | 1:1000 |
| H4, monoclonal | Abcam | mouse | ab31830 |  | 1:500 |

**Table 5.**
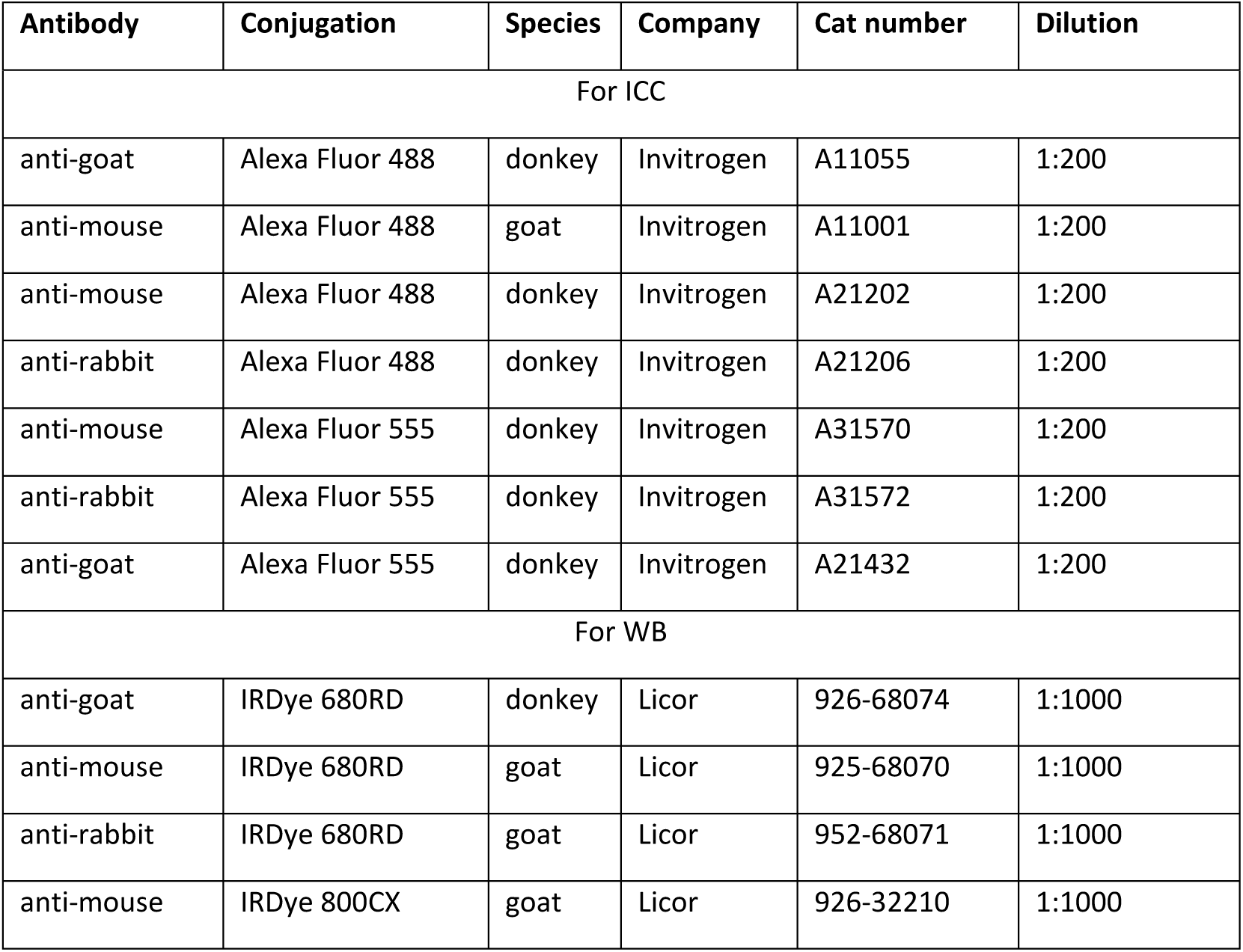
Secondary antibodies used for immunocytochemistry and western blots.

| Antibody | Conjugation | Species | Company | Cat number | Dilution |
| --- | --- | --- | --- | --- | --- |

| For ICC |  |  |  |  |  |
| --- | --- | --- | --- | --- | --- |
| anti-goat | Alexa Fluor 488 | donkey | Invitrogen | A11055 | 1:200 |
| anti-mouse | Alexa Fluor 488 | goat | Invitrogen | A11001 | 1:200 |
| anti-mouse | Alexa Fluor 488 | donkey | Invitrogen | A21202 | 1:200 |
| anti-rabbit | Alexa Fluor 488 | donkey | Invitrogen | A21206 | 1:200 |
| anti-mouse | Alexa Fluor 555 | donkey | Invitrogen | A31570 | 1:200 |
| anti-rabbit | Alexa Fluor 555 | donkey | Invitrogen | A31572 | 1:200 |
| anti-goat | Alexa Fluor 555 | donkey | Invitrogen | A21432 | 1:200 |
| For WB |  |  |  |  |  |
| anti-goat | IRDye 680RD | donkey | Licor | 926-68074 | 1:1000 |
| anti-mouse | IRDye 680RD | goat | Licor | 925-68070 | 1:1000 |
| anti-rabbit | IRDye 680RD | goat | Licor | 952-68071 | 1:1000 |
| anti-mouse | IRDye 800CX | goat | Licor | 926-32210 | 1:1000 |

### Signal quantification

Pre-processing of images was performed by segmentation of single nuclei to feed into a macro for signal quantification. This step was performed using another ImageJ macro which takes the raw Z-stack image and asks the user to select a Z-plane from which to isolate nuclei. This is to avoid nuclei and chromocenters from different planes overlapping. The macro then selects the DAPI channel and uses the ‘Default’ thresholding method to find all the nuclei and separates nuclei which are touching using the ‘Watershed’ function. The user then manually removes any nuclei which have been incorrectly segmented with the ‘Watershed’ function. The remaining nuclei are then added to the Region of Interest (ROI) manager. The ‘Enlarge’ function is used to create a buffer region around the individual nuclei, which are then saved as 16-bit tiffs and used for the following macro (n>100 nuclei). The quantification of signal distribution to the chromocenters was performed through an ImageJ macro which uses the ‘Intermodes’ threshold to define the area of the nucleus in the DAPI channel as a ROI. Nuclei touching the edge are excluded and the ‘Watershed’ feature separates touching nuclei. Automatic thresholding using the ‘Default’ method is then performed with a bottom cut-off equal to 1.2 standard deviations + mean signal intensity to determine the area and position of the chromocenters as ROIs. Nuclei must have at least 1 chromocenter within the thresholded signal to proceed through the macro. The combined area of the chromocenters is then subtracted from that of the nucleus to give the area of the nucleoplasm – also added as ROI. The mean signal intensity for all ROIs in all channels is then measured. The signal intensity at the chromocenters for each channel of interest is then calculated as the intensity at chromocenter divided by the intensity in the nucleoplasm, multiplied by 100 (Supp fig.1A).

To calculate the percentage of the chromocenter occupied by increased signal of interest, in the channel(s) of interest, the area of the nucleus is thresholded using the ‘Intermodes’ method as an ROI. Automatic thresholding using the ‘Default’ method is then performed with a bottom cut-off equal to 1.2 standard deviations + mean signal intensity to determine the area and position of the foci in the channel as ROIs. The area of the foci is then divided by that of the chromocenter, defined in the DAPI channel and multiplied by 100 to give the percentage of the chromocenter occupied by increased signal.

### Live cell imaging

Cells for live-cell imaging were plated on 0.2% gelatin-coated glass bottomed microslides (Ibidi, #80826) and left to settle overnight at 37°C and 5% CO_2_. Slides were then placed in a temperature regulated imaging chamber on a spinning disc confocal microscope (Dragonfly, Andor) Images were obtained using a 40x air objective and captured with iXon camera. For each condition a Z-stack was captured every 3 minutes for 2 hours.

### Protein extraction and quantification

Snap frozen cell pellets were resuspended in RIPA buffer (Thermofisher, #89901) with phosphatase inhibitor at 1X (Thermofisher (100X) Cat# 1862495) and cOmpleteTM protease inhibitor at 1X (Sigma (7X) # 11697498001) until the pellet is completely dissolved. The samples were left on ice for 10 min, after which 1µl of Benzonase (Sigma, #E1014-5KU) was added and left on ice for 30 min to digest the DNA. The samples were then sonicated (method: on/off 30sec/30sec, 15 cycles, on High) (Diagenode) and centrifuged for 10 min at 4°C at 14 000 rpm and the supernatant transferred to a fresh tube. Quantification of protein concentration was performed using Pierce™ BCA protein assay kit (Thermofisher, #23225) following manufacturer’s protocol. Standards were diluted as per provided instructions in the kit.

### Western blot

15µg of protein per sample was incubated with 1X reducing agent (Thermofisher (10X), #NP0004) and 1X LDS sample buffer (Thermofisher (4X), #NP0007) at 100°C for 10min. The samples were then briefly centrifuged and loaded on a precast NuPAGE™ 4-12% Bis-Tris Protein Gel, 1.0 mm (Thermofisher, #NP032) and run in 1X MES SDS running buffer (Thermofisher, #NP0002) with 1:400 concentration of antioxidant (Thermofisher, #NP0005) added to gel compartment of tank. The gel was run at 90V for 1.5h. The manufacturer’s protocol (P0) for iBlot2 Dry blotting was adapted for transfer of larger proteins such as DNMT1 by extending the 20V step to 8 min. Following transfer, the membrane was incubated with Odyssey Blocking Buffer (LI-COR, #927-50000) for 1h on a rocker at RT. Appropriate primary antibodies (Table 2.4) were diluted in 50/50 solution of 1% Tween PBS (TPBS) and Odyssey Blocking Buffer and added to the membranes to incubate O/N on rocker at 4°C in opaque container. The membrane was then washed 3×10 min with TPBS at RT on a rocker after which the membrane was incubated with the secondary antibody (Table 2.5) diluted at 1:200 in the same 50/50 solution for 1h at RT on rocker. After 3×10 min washes with TPBS the membrane was stored in PBS at 4°C until visualisation using a LI-COR Odyssey FC imaging system.

### DNA extraction and quantification

Snap frozen pellets were resuspended in 0.5ml TE buffer per sample (100mM Tris, pH 8.5, 5mM EDTA, 0.2% SDS, 200mM NaCl). 2µl RNAse cocktail was added to each tube and left for 1h at 37°C at 500rpm in a thermoshaker to degrade the RNA in the samples. 20µl of proteinase K was added to each tube and left on the thermoshaker overnight at 56°C to digest proteins. The next day, an equal volume of room temperature UltraPure™ Phenol: Chloroform: Isoamyl Alcohol (#15593-031) was added to each sample inside a chemical hood, mixed by pipetting, and centrifuged at 16 000rpm for 8 min to separate the phases. The upper aqueous phase which contains the DNA was collected into a new tube and an equal volume of phenol was added and mixed and the spinning step was repeated. After collecting the aqueous phase into a new tube, an equal volume of chloroform was added and mixed by pipetting and spun down for 4 minutes at 16 000rpm. This step was repeated twice. 3M sodium acetate was added at 1/10^th^ of solution volume, followed by 100% Ethanol at 2.5 times the solution volume to precipitate the DNA. Samples were left for a minimum of 1h at −20°C to facilitate precipitation. The samples were then spun down to pellet the DNA at 16000rpm for 8 minutes. The pellet was washed twice with 75% ethanol and left to air dry in a chemical fume hood for 1 min. The DNA was resuspended in the appropriate amount of TE buffer (without SDS) and quantified using the Qubit dsDNA BR assay kit (Q32850).

### Probe labelling for Southern Blot

Plasmids containing minor satellite fragments (kindly donated by Shelagh Boyle, Bickmore lab) were used in reaction to generate Digoxigening-11-dUTP (DIG-UTP) labelled probes in 2µl of 10x Nick Translation salts (0.5M Tris pH7.5, 0.1M MgS04, 1mM DTT, 0.5mg/ml BSA fraction V (Sigma)), 2.5µl each of 0.5mM dATP, dCTP and dGTP, 1µl 0.5mM dTTP, 1.5µl DIG-UTP, 1µl of diluted in ice-cold water (1 in 5) DNase I (Roche), 1µl DNA polymerase 1 (Invitrogen, #18010-017) and 500ng of plasmid DNA and MilliQ water up to total reaction volume of 20µl. The reaction mix was incubated at 16°C for 90 minutes and was inactivated with 2µl 20% SDS and 3µl 0.5mM EDTA (pH 8) and made up to 90µl with 65µl TE. The sample was put through a Quick spin column (Roche cat#11273973001) as per manufacturer’s instructions.

### Southern blot

100ng of gDNA of each sample was digested with methylation-sensitive restriction enzyme HpyCH4IV (also known as MaeII, NEB, R0619S) overnight at 37°C. Samples were loaded on a 1% agarose 0.5X TBE gel along with DIG-labelled DNA molecular weight ladder (Roche, #11218590910), without ethidium bromide (EtBr), and run for approx. 3h at 115V. The gel was later submerged in denaturation solution (0.5M NaOH, 1.5M NaCl) twice for 15 min each, rocking at RT. The gel was rinsed with MilliQ water and neutralized in neutralising solution (0.5M Tris pH 7.5, 1.5M NaCl) twice for 15 min each, on the rocker at RT, then equilibrated in 20X SSC for 10 min, rocking. The stack for overnight transfer of the DNA to the membrane was assembled in 20x SSC - using 4x 3MM Whatmann paper wicks and Hybond N+ pre-soaked membrane. The transferred DNA was fixed on the membrane by UV irradiation crosslinking using the Stratalinker 1500 at 0.15J for 30 sec. The membrane was rinsed with MilliQ water and pre-hybridised with DIG Easy Hyb buffer from DIG-High Prime DNA Labelling and Detection Starter Kit II (Roche, #11585614910) for 30 min at 55°C rocking in EnviroGenie. The DIG-labelled probe was denatured at 95°C for 5 minutes and quickly cooled on ice before adding to the appropriate amount of DIG Easy Hyb buffer. Next, the pre-hyb buffer was removed and the membrane was left in the probe-containing buffer to hybridise for minimum of 4 hours at 55°C rocking in EnviroGenie. The probe-containing buffer was reused up to 5 times. The membrane was then washed with pre-heated to 65°C low stringency buffer (2x SSC, 0.1% SDS) twice for 5 minutes each at RT with rocking, and twice for 15 minutes each with High stringency buffer (0.5x SSC, 0.1% SDS) at 65°C with rocking in EnviroGenie to remove unbound probe. The following steps were all performed at room temperature on the rocker. The membrane was equilibrated by washing with Maleic Acid Buffer with Tween (MABT, 0.1M Maleic Acid, 0.15M NaCl, pH adjusted to 7.5, 0.3% Tween 20) for 5 min, then incubated with 1x blocking solution from kit diluted in Maleic Acid Buffer (0.1M Maleic Acid, 0.15M NaCl, pH adjusted to 7.5), followed by diluted blocking solution with anti-DIG antibody from kit at 1:1000 each of the steps lasting 30 minutes. The membrane was then washed twice with MABT for 15 min each and equilibrated with detection buffer (0.1M Tris-HCl, 0.1M NaCl, pH 9.5) for 5 min. CSPD solution (Roche, #11755633001) was used to produce Chemiluminescent reaction which was visualised on X-Ray film or on ImageQuant visualizer.

### RNA extraction and RNA sequencing

Total RNA was extracted using the Direct-Zol RNA extraction kit (Zymo Research, RN2051) and quantified using the Qubit HS RNA assay kit. RNA integrity was assessed by RNA Nano chip on Agilent 2100 Bioanalyzer and RNA samples with RIN score 9 or higher were used for library preparation. First, 500ng of total RNA was enriched for Poly(A) mRNA by using Oligo(dT)25 magnetic beads (NEB, E7490S). Next, libraries were prepared using the NEBNext® Ultra™ II Directional RNA Library Prep Kit for Illumina® (NEB, E7765) following the standard protocol and using 10 cycles for PCR amplification. Quality of the libraries was assessed using the High Sensitivity DNA chip on Agilent 2100 Bioanalyzer and submitted for paired-end 75bp sequencing on NextSeq 550, carried out at the Edinburgh Wellcome Trust Clinical Research Facility, Western General Hospital, Edinburgh, UK. Sequencing data is available at the GEO repository: GSE246241.

### Quantification of repeat expression from RNA sequencing data

Adaptors were removed from RNA-seq reads using TrimGalore! 0.4.1 (paired end, illumina, stringency 5), and aligned to the mm10 mouse genome using TopHat 2.1.0 (very sensitive, inner distance −1 ± 58, no coverage search, max multihits 1). Read co-ordinates were intersected with UCSC genome browser RepeatMasker track co-ordinates using BEDTools 2.25.0, filtered to ensure that each pair or singleton was assigned to only one repeat location, and the number of sequences belonging to each type of repeat summed. Repeats that were expressed at a low level (<5 fpm in more than half the samples) were removed then those belonging to LTR, LINE and SINE Repeatmasker classes that were significantly upregulated (FDR< 0.05, logFC > 0) were identified using the DESeq2 package in R.

### De novo analysis of published ChIP data

Single-end ChIP-seq reads were downloaded from NCBI Geo database (GSM4024115, GSM4024116, GSM4024117). Raw reads were subjected to adaptor trimming and quality filtering using fastp (version 0.23.2) with default parameters. Reads were aligned to mouse genome (UCSC, mm10 assembly) with bowtie2 (version 2.2.6) in two ways: with default parameters so that a single best alignment is reported for each read (-k=1) and with up to 10 of the top alignments reported for the multi-mapping reads (-k=10). This was to make sure that observation is not affected by multi-mapping of the reads because of repeat nature of target sequences in question. Alignments were converted to bam format using samtools (Version: 1.3) and sorted by coordinates and PCR duplicates were removed with samtools (rmdup -s). Coordinates for Satellite repeats were extracted from RepeatMasker track hosted at UCSC genome browser for mm10 assembly of the mouse genome. Then we used bedtools (v2.25.0) to count reads that overlap with satellite repeats (using intersect -c) with or without removing PCR duplicates. Read counts were expressed as percentage of total reads that overlap satellite repeats in the ChIP library for comparison.

### 5mC HPLC Mass spectrometry

1-2 µg of DNA was extracted using Phenol Chloroform extraction method described above. Samples were sent to the Kriaucionis group in Oxford where mass spectrometry analysis was performed using their previously published protocols (34). Samples were injected three times for technical replicates. Results were returned as total number for each base and for each type of base modification (mC and hmC). The quantification bar plot was generated in GraphPrism9 as the percentage of 5mC to total guanines.

## Results

### Polycomb complexes are redirected to PCH following loss of DNA methylation in mESCs

Previous work has shown that the Polycomb homologue and PRC1 component, CBX7, accumulates on the facultative inactive X-chromosome in female XX mammalian cells (35). It is hypothesised that PRC1 is directed there by CBX7 chromodomain which recognises H3K27me3. To test this, we used a published H3K27me3 reader reporter containing a double *Cbx7* chromodomain fused to GFP (Fig 1A). This reporter has previously been shown to specifically bind to H3K27me3 by chromatin immunoprecipitation (24). We introduced the piggyBac *Cbx7*-GFP reporter into normally methylated mESCs (WT J1) and hypomethylated *Dnmt1*^−/−^ *Dnmt3a*^−/−^ *Dnmt3b*^−/−^ triple KO (TKO) cells (Figure 1). H3K27me3 has previously been shown to accumulate at PCH in hypomethylated TKO mESCs by immunostaining (19, 21). As expected, in J1 cells (normal methylation) the reporter shows a similar nuclear distribution pattern as observed by Villaseñor et al (2020)(24) (Fig 1B, top panel), while in the hypomethylated TKO cells the reporter displays a clustering punctate pattern, reminiscent of heterochromatic foci, as predicted (Fig 1B bottom panel). Immunocytochemistry (ICC) microscopy on TKO fixed cells showed that the *Cbx7*-GFP foci overlap with the DAPI-defined PCH, and co-stained for the PRC1 mark, H2AK119ub1, confirming the redistribution of both PcG marks in the same cell Fig 1C). In contrast, both the reporter and the H2AK119ub1 antibody showed dispersed signal throughout the nucleus of J1 cells, corresponding to the live-cell imaging result. We then used a custom designed ImageJ macro to quantify in an unbiased manner the signal within the DAPI foci for the reporter and the antibody. The quantification showed that both *Cbx7*-GFP and H2AK119ub1 signals are significantly increased by 1.5-2.0 fold within the hypomethylated chromocenters of TKO cells compared to normally methylated chromocenters in J1s (Fig 1D). This is also supported by a higher percentage of the chromocenter area being occupied by increased *Cbx7*-GFP reporter and H2AK119ub1 antibody signals in the TKOs (Supp fig 1B). These results support previous published reports suggesting that both PRC1 and PRC2 relocalise to PCH in the absence of DNA methylation (19, 21, 22, 36). Furthermore, H2AK119ub1 signal intensity is significantly increased within *Cbx7*-GFP defined foci relative to the nucleoplasm in TKO but not J1 cells, whereas H3K9me3 signal intensity is not (Fig 1E). *Cbx7*-GFP signal similarly significantly overlaps with H2AK119ub1 but not H3K9me3 specifically in hypomethylated TKO cells (Supp fig 1C). These data suggest the *Cbx7*-GFP reader is not recognising the H3K9me3 mark which occupies the PCH in both J1 and TKO cells (19), but is colocalising with Polycomb-mediated chromatin modifications that accumulate at PCH in response to DNA hypomethylation.

**Fig. 1.**
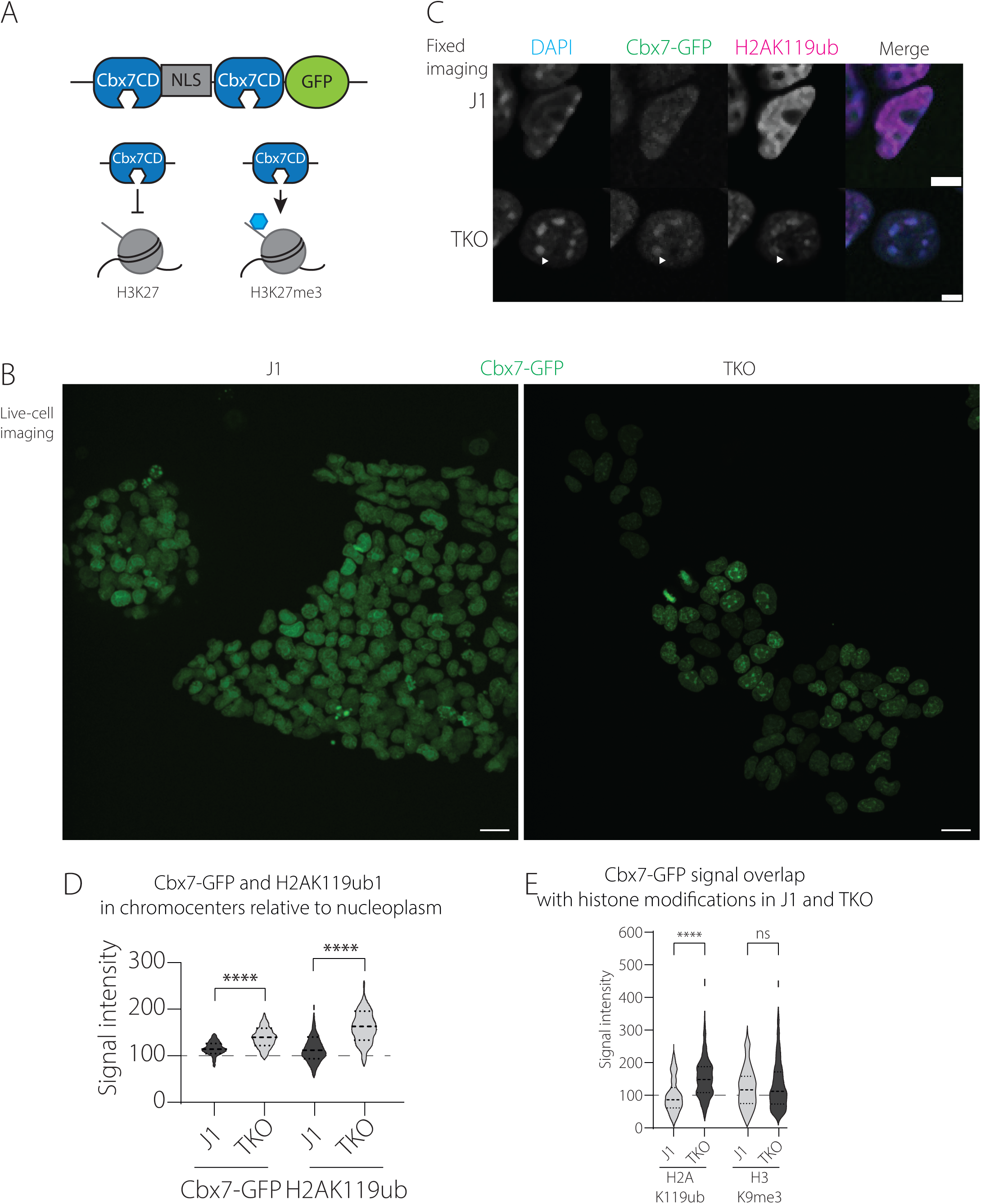
Cbx7-GFP detects H3K27me3 and forms clusters over PCH. (A) Graphical representation of the Cbx7-GFP reporter. The reporter contains two Cbx7 chromodomains (Cbx7CD) flanking a Nuclear Localisation Signal (NLS) sequence followed by a Green Fluorescent Protein domain. The reporter was generated in the Baubec lab to recognise specifically tri-methylated H3K27 (right) and not unmethylated H3K27 (left). We integrated the reporter sequence into a piggybac vector for this study. (B) Single timepoint snapshots from maximum intensity z-projection of J1 and TKO live cells with the Cbx7-GFP reporter (green signal), imaged over 2 hours. Scale bar corresponds to 20µm. (C) IF on J1 and TKO cells containing Cbx7-GFP reporter stained for H2AK119ub counterstained with DAPI. Representative nuclei from single slice of 3D stack are shown. Z-stacks obtained using 40x air objective. Scale bar corresponds to 5µm. Arrowheads point to a representative chromocenter at which all three signals overlap in the TKO cells. (D) Violin plot showing mean signal intensity of Cbx7-GFP reporter/H2AK119ub antibody signal at chromocenter normalised to that in the nucleoplasm in J1 and TKO cells. Grey dashed line corresponds the threshold above which the signal is considered increased at the chromocenter compared to the nucleoplasm. Significance was determined by Kruskal-Wallis (one-way Anova) test where **** p≤0.0001. (E) Violin plot of mean signal intensity of antibody (H2AK119ub and H3K9me3) signal at Cbx7-GFP-defined foci normalised to that in the nucleoplasm in J1 and TKO cells. Grey dashed line corresponds the threshold above which the signal is considered increased at the Cbx7-GFP locus compared to the nucleoplasm. Significance was determined by Kruskal-Wallis (one-way Anova) test with FDR correction-Benjamini and Hochberg where **** p≤0.0001. Test was performed between J1 and TKO samples for each antibody.

Established knockout mESC lines may acquire adaptive changes in response to selective pressure to stabilise their hypomethylated heterochromatin near the centromeres, to protect genome integrity. In contrast, the gradual loss of DNA methylation is more comparable to the DNA demethylation programming that occurs in early preimplantation embryos, which arises through active and passive (replicative) DNA demethylation in the presence of transient de novo methylation (37). To recapitulate a dynamic loss of DNA methylation, we used a previously published *Dnmt1* tetOFF (*Dnmt1*^tet/tet^) ES cell line (Fig 2A) (29). In this system, doxycycline (dox) treatment suppresses *Dnmt1* transcription, resulting in downregulation of the endogenous DNMT1 protein (Fig 2B). This is concomitant with DNA hypomethylation as evidenced by southern blotting for a normally methylated minor satellite region resident at PCH and by HPLC assays for global 5mC levels in genomic DNA (Fig 2 C and D). By day 4 of dox treatment, the minor satellites are fully hypomethylated and 5mC levels have reduced by 50%. Western blotting shows that global levels of H3K27me3, H2AK119ub1 and H3K9me3 levels are unaltered during the 7-day dox treatment, suggesting that any change in localisation is not due to a change in global levels of these marks (Fig 2E).

**Fig. 2.**
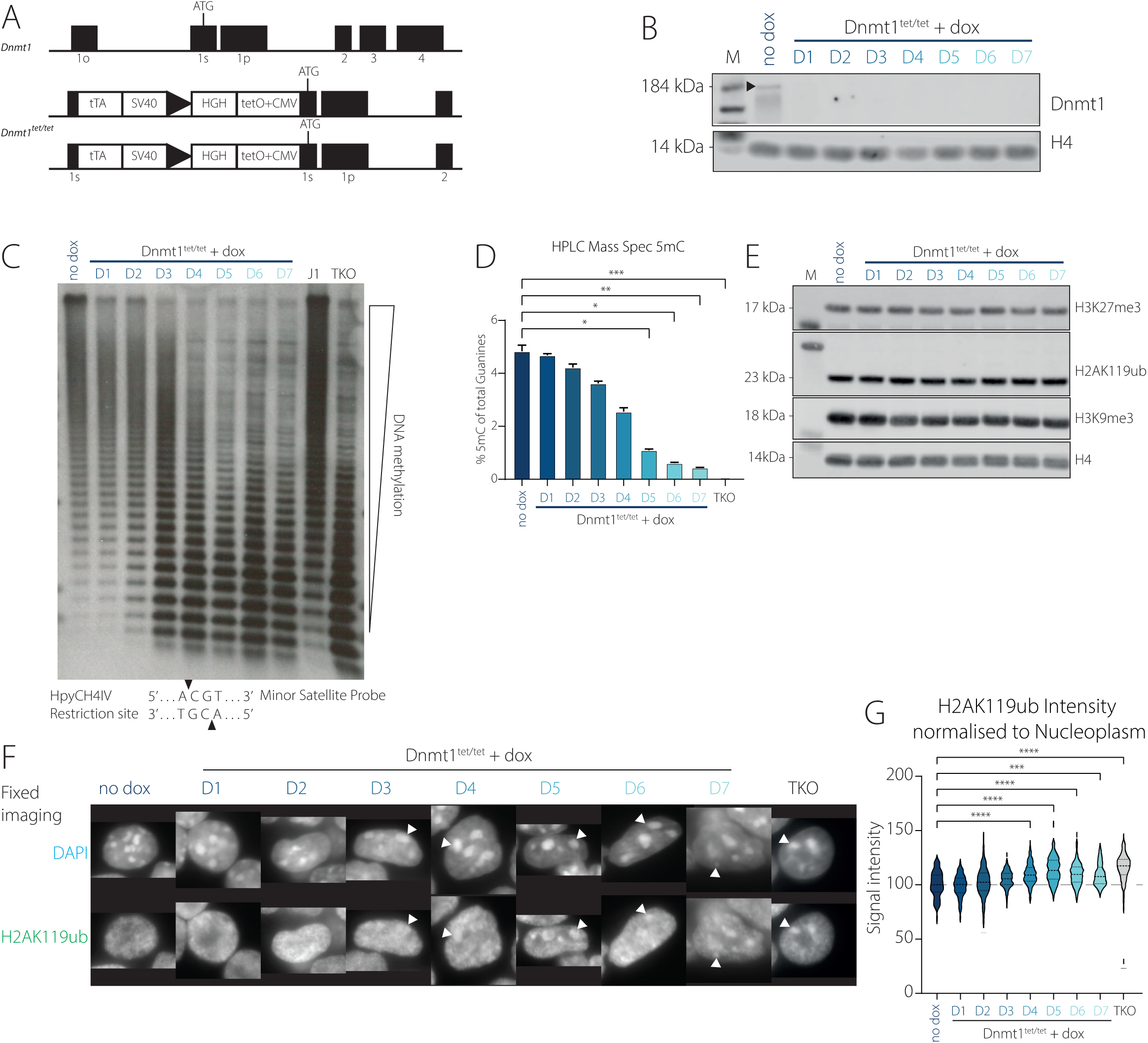
PRC1 relocalises to PCH as the locus hypomethylates. (A) Graphical representation of the Dnmt1^tet/tet^ cell line. Donated by Chaillet lab. Modified from Borowczyk et al., 2009 – lacking neomycin and puromycin resistance cassettes. (B) Western blot for Dnmt1 and H4 (loading control) proteins over 7 day dox treatment in for Dnmt1^tet/tet^ cells. M – Molecular ladder. Arrow points to expected band for Dnmt1 protein. (C) Southern blot using methylation sensitive restriction digest (HpyCH4IV enzyme –cut site shown below) followed by hybridisation using probe for minor satellite repeats in Dnmt1^tet/tet^ cells with (over 7 days) or without dox and in J1 and TKO cells. Methylated DNA cannot be digested and remains as high MW bands. Unmethylated DNA is digested and separates into polymers of satellite repeats. (D) HPLC Mass spectrometry analysis of global 5mC levels as percentage of total Guanines over 7 day dox treatment in for Dnmt1^tet/tet^ cells and TKO cells as control. Significance was determined by Kruskal-Wallis (one-way Anova) test with FDR correction – Benjamini, Krieger and Yekutieli. (E) Western blot for H3K27me3, H2AK119ub, H3K9me3 and H4 (loading control) over 7 day dox treatment in for Dnmt1^tet/tet^ cells. M – Molecular ladder. (F) Fixed immunofluorescence (IF) in Dnmt1^tet/tet^ cells over 7 days dox treatment and TKO cells stained for H2AK119ub (bottom row) and counterstained for DAPI (top row). Representative nuclei from single slice of 3D stack are shown. Z-stacks obtained using 40x air objective on an epifluorescent microscope. Scale bar corresponds to 5µm. Arrowheads point to a representative chromocenter at which the DAPI and antibody signals overlap. (G) Violin plot showing mean signal intensity of H2AK119ub antibody signal at chromocenter normalised to that in the nucleoplasm in Dnmt1^tet/tet^ cells over 7 days dox treatment and TKO cells. Grey dashed line corresponds the threshold above which the signal is considered increased at the chromocenter compared to the nucleoplasm. Significance was determined by Kruskal-Wallis (one-way Anova) test where **** p≤0.0001.

We then aimed to determine the timeframe relative to the hypomethylation of the satellites in which H2AK119ub1 relocalises to the PCH. We stained *Dnmt1*^tet/tet^ cells over the course of a 7 day dox treatment for H2AK119ub1 and found that the H2AK119ub1 signal aggregates at DAPI-rich foci as early as day 3-4 of dox treatment (Fig 2F). Quantification of the H2AK119ub1 signal at the chromocenters confirmed that the PRC1 modification is significantly enriched at day 4 of dox treatment, coinciding with the hypomethylation of the region (Fig 2G).

Having established that the Cbx7 live-cell reporter can track H3K27me3 localisation, we integrated the reporter into the *Dnmt1^tet^*^/tet^ cell line. By day 4 of dox treatment aggregate *Cbx7*-GFP foci form, which are stable through interphase (Fig 3A and B, top panel) The *Cbx7*-GFP signal is associated with mitotic chromosomes that re-form punctate foci at the end of mitosis, suggesting that the reporter is able to follow the modification through different chromatin conformations. In this system, live-cell imaging with CBX7 presents a quick and direct readout of the relocalisation of H3K27me3 and throughout the cell cycle. Co-staining the *Cbx7*-GFP reporter with H2AK119ub1 antibody in fixed cells showed aggregation into foci at day 4 of dox-treated cells compared to control, similar to TKO cells (Fig 3C) and this redistribution appeared to be reversible upon dox withdrawal (Fig 3D). This is supported by a significant increase of the signal inside the chromocenters compared to the nucleoplasm in dox treatment for *both* Cbx7-GFP and H2AK119ub1 (Fig 3E and Supp fig 1D).

**Fig. 3.**
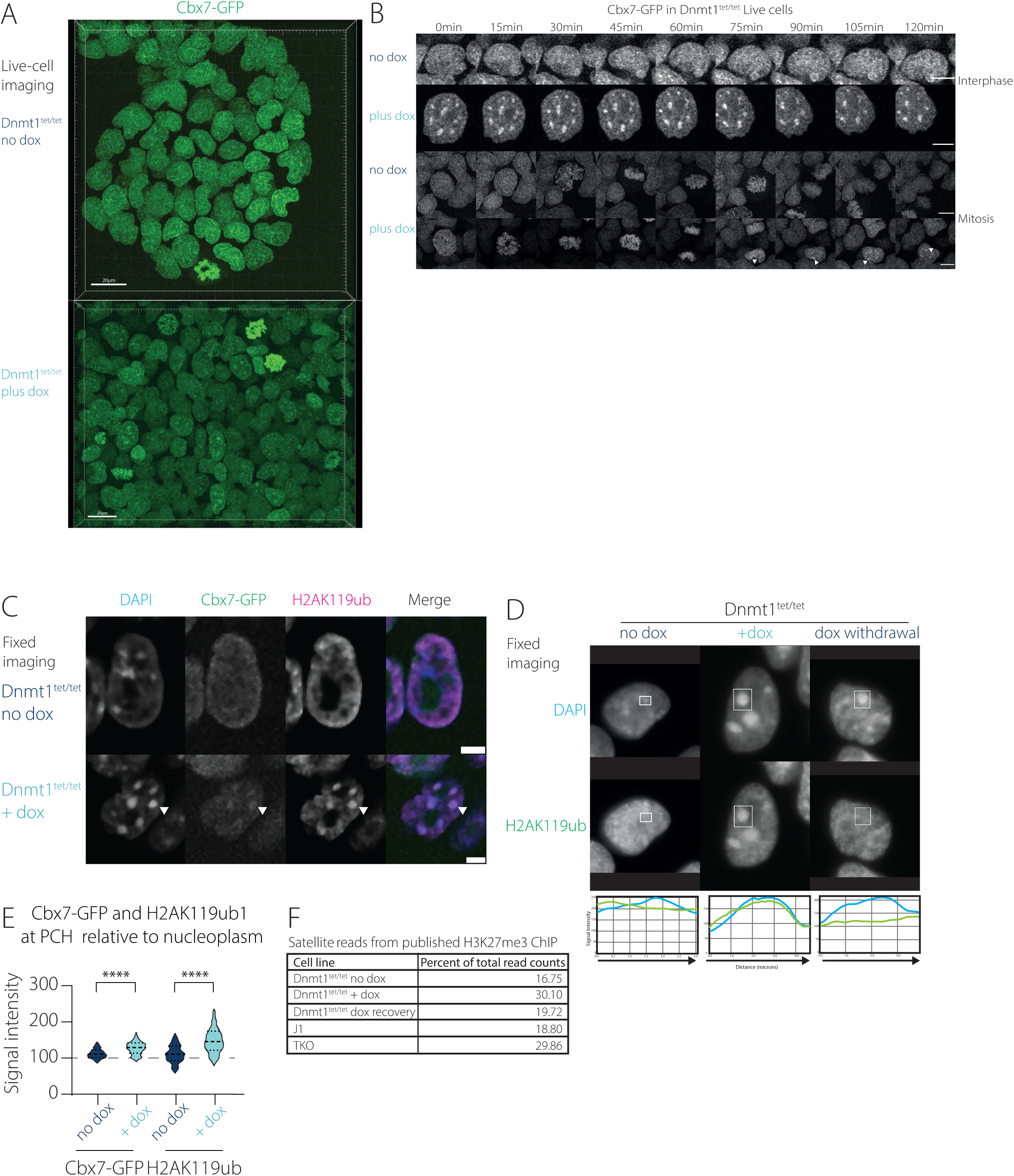
Polycomb complexes are redirected to PCH following loss of DNA methylation in mESC. (A) Single timepoint snapshots from maximum intensity z-projection of Dnmt1^tet/tet^ cells without (top panel) or with dox for 4 days (bottom panel) with the Cbx7-GFP reporter (green signal), imaged over 2 hours. Scale bar corresponds to 20µm. (B) Live-cell imaging of Dnmt1^tet/tet^ cells without or with dox for 4 days containing the Cbx7-GFP reporter, imaged over 2 hours, image taken every 3 min. Representative nuclei at interphase (top two panels) and mitosis (bottom two panels) shown at indicated time points (every 15min). Scale bar corresponds to 10µm. Arrowheads point to clustering of Cbx7-GFP signal under dox conditions following exit from mitosis. (C) IF on Dnmt1^tet/tet^ cells containing Cbx7-GFP reporter stained for H2AK119ub counterstained with DAPI. Representative nuclei from single slice of 3D stack are shown. Z-stacks obtained using 40x air objective. Scale bar corresponds to 5µm. Arrowheads point to a representative chromocenter at which all three signals overlap. (D) IF for H2AK119ub counterstained with DAPI in Dnmt1^tet/tet^ cells in presence of dox and following dox recovery. Representative nuclei from single slice of 3D stack are shown. Z-stacks obtained using 40x air objective on an epifluorescent microscope. Scale bar corresponds to 5µm. Plot shows the signal intensity of each of the channels along the diagonal of the rectangle in each channel at the specific chromocenter (blue line - DAPI channel, green line - antibody signal. (E) Violin plot showing mean signal intensity of Cbx7-GFP reporter/H2AK119ub antibody signal at chromocenter normalised to that in the nucleoplasm in Dnmt1^tet/tet^ cells with or without dox. Grey dashed line corresponds the threshold above which the signal is considered increased at the chromocenter compared to the nucleoplasm. Significance was determined by Kruskal-Wallis (one-way Anova) test where **** p≤0.0001. Dark blue signifies high DNA methylation, Light blue corresponds to low DNA methylation. (F) Percentage of reads corresponding to satellite repeats to total number of reads from published H3K27me3 ChIP data on Dnmt1^tet/tet^ cells without, with and upon dox recovery and in J1 and TKO.

Our imaging of the *Cbx7*-GFP reporter suggests that Polycomb-dependent chromatin modifications become enriched at PCH upon depletion of DNMT1. To verify this observation, we analysed the abundance of satellite DNA in our previously published H3K27me3 ChIP data from *Dnmt1*^tet/tet^ cells in the presence or absence of dox and upon dox recovery. Consistent with the imaging data and previous results in other hypomethylated contexts (21) we found more H3K27me3-enriched satellite DNA in hypomethylated conditions (dox; TKO) than normally methylated conditions (no dox; J1) and upon recovery of DNA methylation (dox recovery) (Fig 3F). Thus, the enrichment of Polycomb-dependent modifications at PCH in hypomethylated ES cells that we detect by imaging likely reflects an increase in association of Polycomb-dependent modifications with satellite chromatin.

The redistribution of Polycomb-dependent modifications to PCH that we have described in interphase cells, is also evident during mitosis. Mitotic chromosomes show an absence of reporter and H2AK119ub staining in centromeric regions in J1 WT cells (Supp fig 1G); however, these regions become stained in TKO cells. These observations are recapitulated in *Dnmt1^tet^*^/tet^ cells where upon addition of dox, the chromosomes obtain a more even distribution of the Polycomb marks that includes the centromere (Supp fig 1H). Taken together, these results suggest that a relocalisation of PRC1 and PRC2 histone modifications to the PCH is contingent with progressive DNA hypomethylation of the region.

The redistribution of Polycomb-dependent modifications to PCH that occurs in response to depletion of DNMT1 is detectable by imaging 4 days after dox treatment (Figure 3). At earlier time-points, the size of the chromocenters becomes larger upon treatment with dox (Supp fig 1E). This change is reversed with longer treatment (Supp fig 1F), suggesting that PCH reorganisation is occurring during the early stages of hypomethylation, which is stabilised following prolonged hypomethylation of the locus.

### PRC1 and PRC2 are recruited to PCH independently of each other

Given that both H3K27me3 and H2AK119ub1 modifications relocalise to the PCH shortly after DNA hypomethylation, we then asked in what order the complexes are relocalising. To that end, we knocked out the catalytic component of PRC2 – EZH2 – in the *Dnmt1*^tet/tet^ background, hereon referred to as *Ezh2* KO (Supp fig 2A-B) and performed a dox treatment. H3K27me3 was not detectable in the *Ezh2* KO (Supp fig 2C), further confirming the role of EZH2 in depositing the trimethylation mark (38). DNMTt1 responds to dox in the *Ezh2* KO cells similarly to the parental cell line (Supp fig 2C). Importantly, the global levels of H2AK119ub1 do not change in the absence of EZH2 or DNMT1 or both (Supp fig 2D). Treatment of both the parental and *Ezh2* KO cell lines with dox resulted in significant H2AK119ub1 relocalisation to PCH in the absence of DNMT1 (Fig 4 A and B). Interestingly, the normalised mean intensity of the ubiquitin mark in the *Ezh2* KO cells with dox was significantly slightly lower than that of the treated parental cell line (Fig 4B). Overall, these results indicate that EZH2 is not necessary for the relocalisation of H2AK119ub1, this modification might have some dependence (either direct or indirect) on the presence of EZH2 (or H3K27me3) at PCH.

**Fig. 4.**
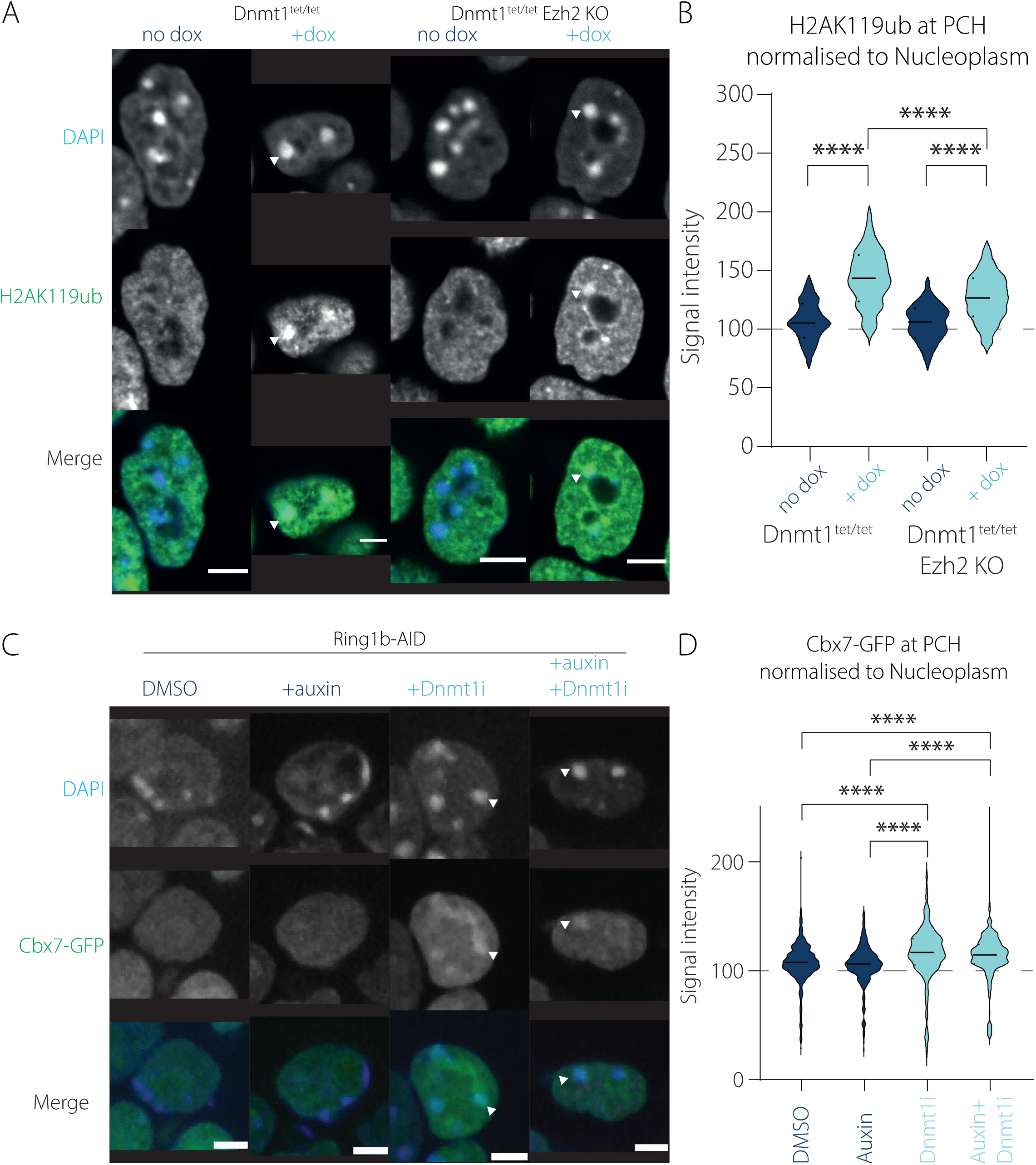
Polycomb complexes are recruited independently to PCH. (A) IF for H2AK119ub counterstained with DAPI in Dnmt1^tet/tet^ cells and Ezh2 KO cells with or without dox. Representative nuclei from single slice of 3D stack are shown. Z-stacks obtained using 40x air objective. Scale bar corresponds to 5µm. Arrowheads point to a representative chromocenter at which the DAPI and antibody signals overlap. (B) Violin plot of mean signal intensity of H2AK119ub antibody signal at chromocenter normalised to that in the nucleoplasm in Dnmt1^tet/tet^ cells and Ezh2 KO cells with or without dox. Grey dashed line corresponds the threshold above which the signal is considered increased at the chromocenter compared to the nucleoplasm. Significance was determined by Kruskal-Wallis (one-way Anova) test where **** p≤0.0001. (C) As in (A) IF for Cbx7-GFP reporter counterstained with DAPI in Ring1b-AID cells treated with DMSO, auxin, Dnmt1i or both auxin and Dnmt1i. Arrowheads point to a representative chromocenter at which the DAPI and reporter signals overlap. (D) As in (B) Violin plot of mean signal intensity of Cbx7-GFP reporter signal at chromocenter normalised to that in the nucleoplasm in Ring1b-AID cells treated with DMSO, auxin, Dnmt1i or both auxin and Dnmt1i.

We also examined the expression of satellite repeats in response to loss of DNA methylation and found no significant change in transcript levels in the absence of *Ezh2* (Supp fig. 2E). This suggests that the relocalisation of H3K27me3 at hypomethylated PCH does not have a supplementary role in repressing satellite transcripts.

To investigate whether H3K27me3 redistribution to PCH relies on PRC1 activity, we used a *Ring1b*-AID cell line where RING1b, a catalytic component of PRC1, is fused to an auxin-inducible degron (AID) (27). Auxin treatment induces the rapid degradation of RING1b protein, which we performed alongside treatment with a recently published inhibitor of DNMT1 activity, GSK-3484862 (termed Dnmt1i), which works by non-covalently inhibiting DNMT1 protein activity (39, 40). Depletion of H2AK119ub1 occurs within 4 days of auxin exposure, however global levels of H3K27me3 and H3K9me3 were not affected (Supp fig 2F). Treatment with 2μM Dnmt1i inhibitor for 4 days did not affect global levels of any of the histone modifications (Supp fig 2F), however, it led to a slight decrease in the DNMT1 protein levels. Similarly to the dox treatment in the *Dnmt1*^tet/tet^ cells, treatment with Dnmt1i led to hypomethylation of minor satellite regions within 4 days of treatment (Supp fig 2G). We introduced the *Cbx7*-GFP reporter in these cells and observed that auxin treatment alone did not induce significant relocalisation of the reporter to the PCH (Fig 4C and D). However, after both Dnmt1i and double treatment with auxin and Dnmt1i, the reporter was displaced and formed characteristic clusters which overlapped with the DAPI foci (Fig 4C). Quantification revealed that the *Cbx7*-GFP reporter signal was significantly increased in both Dnmt1i-treated conditions compared to DMSO or auxin only (Fig 4D), suggesting that the presence of RING1b and subsequently H2AK119ub1 are not necessary for PRC2 recruitment and H3K27me3 deposition at the hypomethylated PCH. Collectively, these observations indicate that PRC1 and PRC2 can be recruited to the hypomethylated PCH independently from each other in mESCs.

### SCML2 localises to PCH irrespective of DNA methylation but does not recruit PcGs to the region

Following the observation that the two PRC activities are recruited independently to the hypomethylated PCH, we then asked which other factors might be involved in their targeting to the region. We hypothesised that a protein capable of this must be expressed in the absence of DNA methylation and it must be able to bind both pericentromeric heterochromatin and PRCs. A potential candidate is Sex comb on midleg-like 2 (SCML2), a germline-specific protein and a variant PRC accessory protein. It has been shown to function as an inhibitor of H2AK119ub1 specifically at PCH and promotes H3K27me3 deposition to the region during meiosis (41). Germline loss of SCML2 results in aberrant facultative heterochromatin formation at the PCH (42). Gene expression analyses have shown that *Scml2* is expressed in hypomethylated ESCs, including TKOs but is not normally expressed in WT mESCs (21). Furthermore, SCML2 has been previously identified as the most enriched protein in a proteomics screen of isolated chromatin segments (PICh) comparing PCH composition between wild-type J1 and hypomethylated TKOs (22).

We performed a macro deletion of *Scml2* (exons 2-26) in the *Dnmt1*^tet/tet^ cell line using CRISPR at the indicated sites (Supp fig 3 A, B and C), referred to as *Scml2* KO. *Dnmt1* expression in *Scml2* KO cells responds to dox treatment similarly to the parental cell line (Supp fig 3D) and methylation-sensitive genes such as *Dazl* become upregulated (Supp fig 3E). We also found significant upregulation of satellite transcripts in the *Scml2* KO in the presence of dox, which was comparable to that of that dox-treated parental line. (Supp fig. 3F).

Imaging of the *Cbx7*-GFP reporter in these cells showed that dox treatment of both the parental and the *Scml2* KO cell lines lead to the significant increase in both *Cbx7*-GFP reporter signal and H2Ak119ub1 aggregation into foci which overlap with the chromocenters compared to their corresponding no dox controls (Fig 5 A-D), suggesting that PRC1 and PRC2 can localise to hypomethylated PCH irrespective of SCML2. Interestingly, the *Cbx7*-GFP reporter signal in the dox-treated *Scml2* KO cells is slightly but significantly lower than that of the parental treated cells, suggesting that SCML2 might facilitate relocalisation or stabilise the H3K27me3 modification at PCH, however the SCML2 protein is not necessary for the process. Importantly, *Scml2* KO dox-treated cells show a significant increase in H2AK119ub1 occupancy of the PCH compared to the parental treated cells. These results suggest that SCML2 is not only unnecessary for PRC1 relocalisation upon hypomethylation of the PCH but might also be preventing the complex’s full occupancy of the region. Taken together, these results indicate that SCML2 does not drive the deposition of either PRC1 or PRC2 to the hypomethylated PCH.

**Fig 5.**
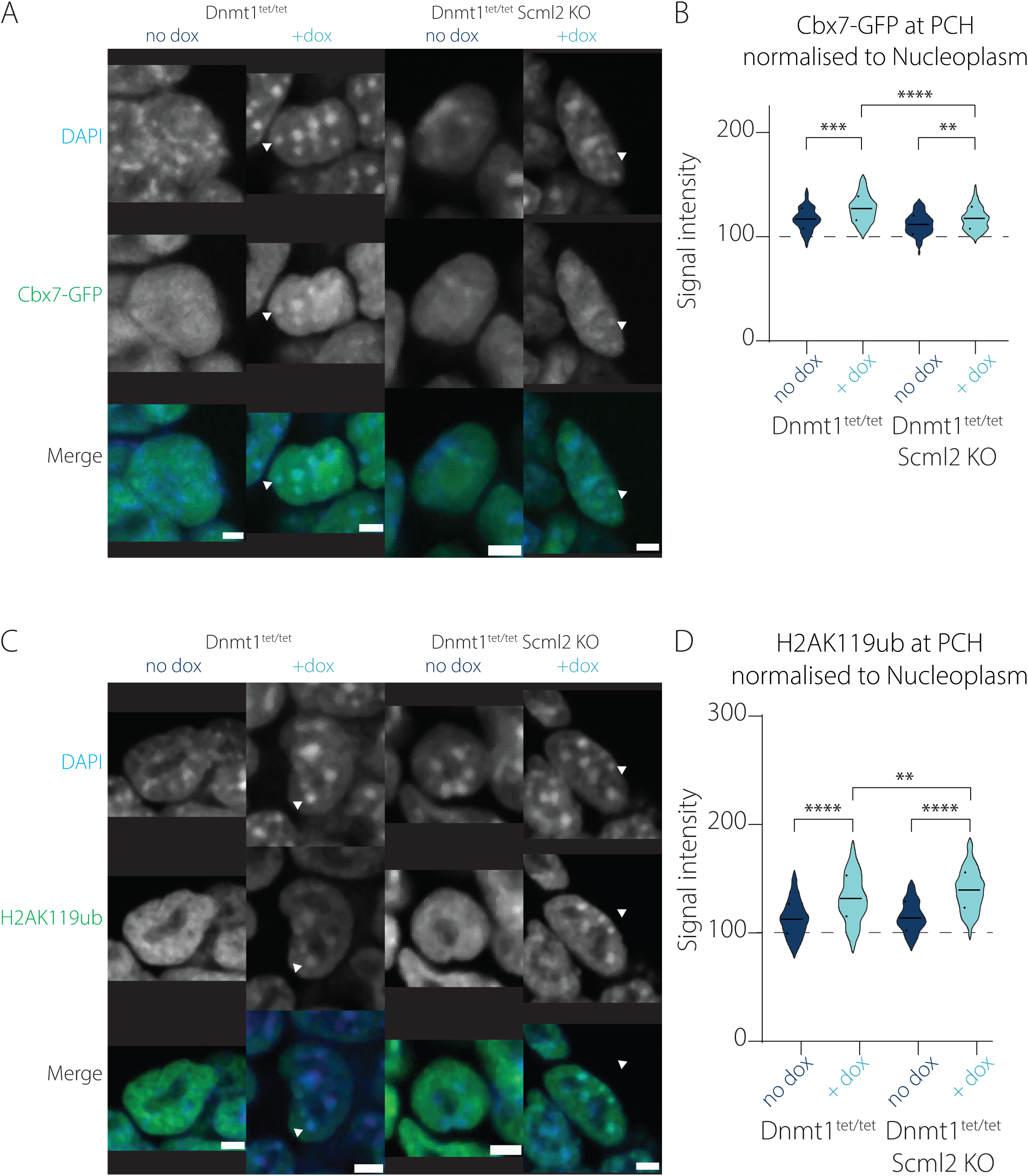
Absence of Scml2 does not prevent PcG relocalisation to hypomethylated PCH. (A) IF for Cbx7-GFP counterstained with DAPI in Dnmt1^tet/tet^ cells and Scml2 KO cells with or without dox. Representative nuclei from single slice of 3D stack are shown. Z-stacks obtained using 40x air objective. Scale bar corresponds to 5µm. Arrowheads point to a representative chromocenter at which the DAPI and reporter signals overlap. (B) Violin plot of mean signal intensity of Cbx7-GFP reporter signal at chromocenter normalised to that in the nucleoplasm in Dnmt1^tet/tet^ cells and Scml2 KO cells with or without dox. Grey dashed line corresponds the threshold above which the signal is considered increased at the chromocenter compared to the nucleoplasm. Significance was determined by Kruskal-Wallis (one-way Anova) test where **** p≤0.0001. Test was performed between every sample. Non-significant samples are not shown. (C) As in (A) IF for H2AK119ub counterstained with DAPI in Dnmt1^tet/tet^ cells and Scml2 KO cells with or without dox. Arrowheads point to a representative chromocenter at which the DAPI and antibody signals overlap. (D) As in (B) Violin plot of mean signal intensity of H2AK119ub antibody signal at chromocenter normalised to that in the nucleoplasm in Dnmt1^tet/tet^ cells and Scml2 KO cells with or without dox.

To test whether SCML2 is sufficient to drive H3K27me3 to PCH in the presence of DNA methylation, we introduced a reporter containing the entire mouse *Scml2* sequence fused to an Enhanced Green Fluorescent Protein (EGFP) at its C-terminal end (*Scml2*-EGFP) (Supp fig 3G) in J1 and TKO cells using the piggyBac system. The *Scml2*-EGFP reporter signal was increased in DAPI-rich foci in both J1 and TKO cells, suggesting that SCML2 can bind the PCH locus independently of DNA methylation (Supp fig 3H). We found no accumulation of H3K27me3 signal in J1 cells expressing *Scml2*-GFP compared to the parental J1 line. These results indicate that the exogenous expression of SCML2 in normally methylated cells is not sufficient to relocalise the H3K27me3 signal to the PCH.

### BEND3 does not contribute to the initial recruitment of either PRC1 or PRC2 in hypomethylated Dnmt1^tet/tet^ ESCs

Following from the observation that the most likely candidate, SCML2, did not appear to be mechanistically necessary for the relocalisation of either of the PcG complexes to the hypomethylated PCH, we then investigated BEND3 as another candidate (22). BEND3 is a transcription regulator during early development known to be important for stabilisation of PRC2 at bivalent genes and CGIs (43, 44). The protein was noted for its enrichment in hypomethylated PCH heterochromatin and was proposed to aid PRC2 recruitment indirectly through its interaction with MBD3/NurD complex (22, 45). The protein was also found as the most highly enriched scoring factor in a PICh assay comparing Suv39h1/2 DKO and WT mESCs and second highest in TKO versus WT cells (22). To test whether BEND3 is responsible for PcG relocalisation to hypomethylated PCH we used CRISPR to knock out the BEND3 gene in the *Dnmt1*^tet/tet^ background (Supp fig 4A, B and C), referred to as *Bend3* KO. Importantly, the *Bend3* KO cell line was responsive to dox similarly to the parental line (Supp fig 4C). *Dazl* expression among other genes (Supp table 1) was significantly upregulated in dox-treated cells (Supp fig 4E) and further significantly upregulated in *Bend3* KO cells in the presence of dox suggesting that *Bend3* prevents full de-repression of *Dnmt1* regulated genes. RNA sequencing confirmed consistent expression of *Bend3* in the parental cells but not in the KO (Supp fig 4C). We also noticed significantly increased transcript levels of satellites in the *Bend3* KO in the presence of dox compared to dox-treated parental cells (Supp fig 4F), suggesting that BEND3 may prevent full activation of satellite expression in hypomethylated *Dnmt1*^tet/tet^ ESCs.

Introducing the *Cbx7*-GFP reporter in the *Bend3* KO cells followed by dox treatment showed significant aggregation of *Cbx7*-GFP and H2AK119ub1 in foci at the chromocenters in both parental and *Bend3* KO dox treated cells compared to the no dox controls (Fig 6A-D). Both the *Cbx7*-GFP reporter and the H2AK119ub1 staining showed more accumulation of signal inside the chromocenters in dox-treated *Bend3* KO compared to the hypomethylated parental cells, suggesting that absence of Bend3 might create a more permissive environment for recruitment of PcGs to PCH. Taken together, these results demonstrate that while BEND3 is not necessary for the initial recruitment of PcGs to PCH in the event of hypomethylation, the protein might be important for maintaining chromatin organisation at the region.

**Fig 6.**
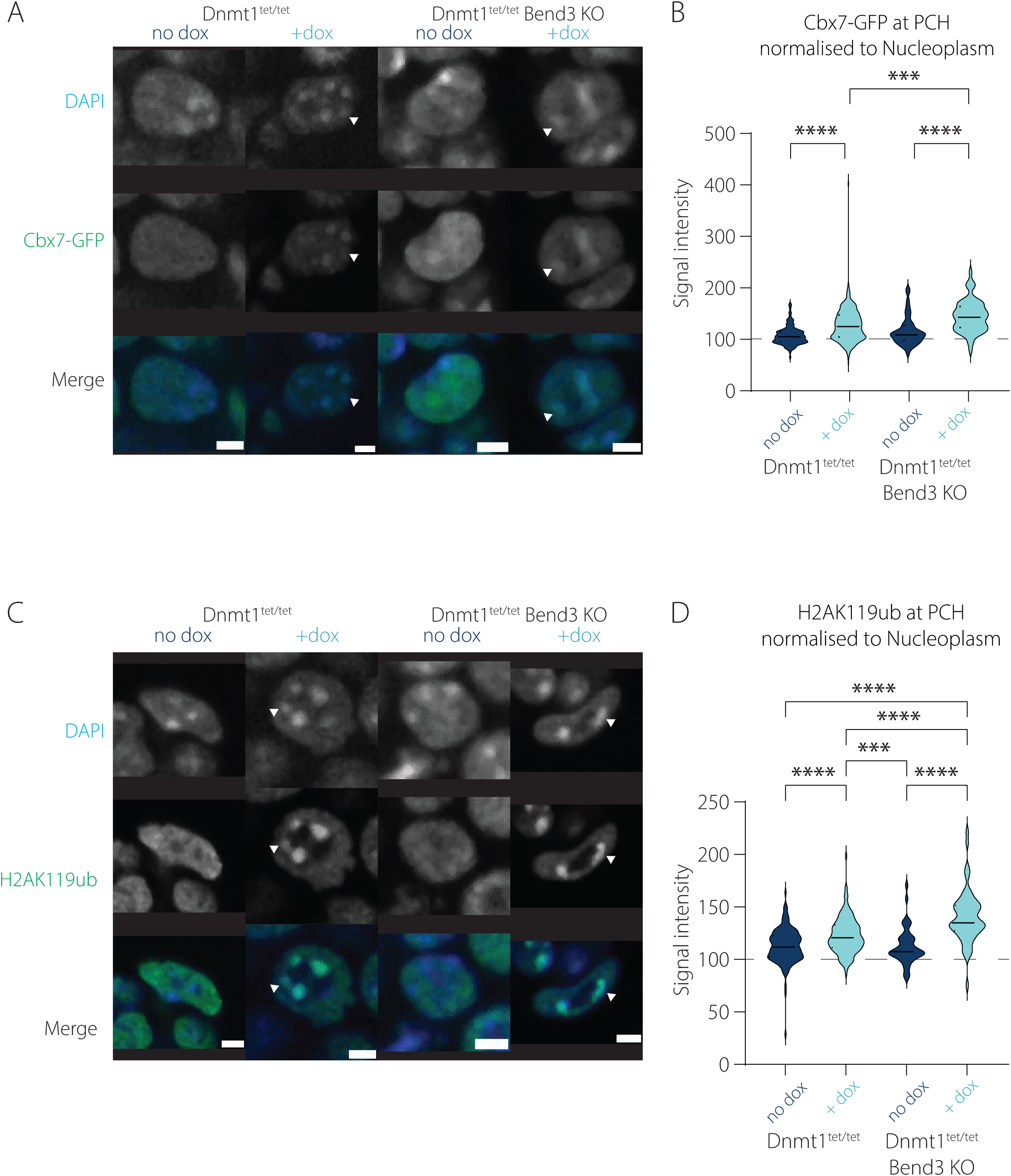
Bend3 is not necessary for the redistribution of PRC1 or PRC2 to PCH. (A) IF for Cbx7-GFP counterstained with DAPI in Dnmt1^tet/tet^ cells and Bend3 KO cells with or without dox. Representative nuclei from single slice of 3D stack are shown. Z-stacks obtained using 40x air objective. Scale bar corresponds to 5µm. Arrowheads point to a representative chromocenter at which the DAPI and reporter signals overlap. (B) Violin plot of mean signal intensity of Cbx7-GFP reporter signal at chromocenter normalised to that in the nucleoplasm in Dnmt1^tet/tet^ cells and Bend3 KO cells with or without dox. Grey dashed line corresponds the threshold above which the signal is considered increased at the chromocenter compared to the nucleoplasm. Significance was determined by Kruskal-Wallis (one-way Anova) test where **** p≤0.0001. Test was performed between every sample. Non-significant samples are not shown. (C) As in (A) IF for H2AK119ub counterstained with DAPI in Dnmt1^tet/tet^ cells and Bend3 KO cells with or without dox. Arrowheads point to a representative chromocenter at which the DAPI and antibody signals overlap. (D) As in (B) Violin plot of mean signal intensity of H2AK119ub antibody signal at chromocenter normalised to that in the nucleoplasm in Dnmt1^tet/tet^ cells and Bend3 KO cells with or without dox.

### KDM2b is not necessary for the recruitment of PcGs to PCH upon hypomethylation

Considering our observations that neither SCML2 nor BEND3 are implicated in recruiting PcGs to hypomethylated PCH, we went on to investigate whether a factor containing a ZF-CxxC domain capable of binding unmethylated CpGs would be sufficient to mediate PcG recruitment. One such factor is KDM2b, an H3K36me1/2 demethylase, which is part of a variant PRC1 complex (46). It recognises non-methylated CpG sites through its ZF-CXXC domain, linking its reported recruitment of PRC1 to CpG hypomethylation (46). KDM2b is highly expressed in mESCs where it occupies almost all CpG island promoters, including bivalent and active genes (19, 46, 47). We used the *Kdm2b*^fl/fl^ cell line derived in the Klose lab in which the ZF-CxxC domain is flanked by LoxP sites allowing for tamoxifen (OHT)-induced excision of this domain (28, 48) (Supp fig 5A). Only treatment with OHT led to the deletion of the region, which did not occur in the Dnmt1i or DMSO treated cells (Supp fig 5B). Dnmt1i treatment for 4 days led to the hypomethylation of the PCH (Supp fig 5C) similarly to that in the *Ring1B*-AID cells (Supp fig 2G).

Upon introduction of the *Cbx7*-GFP reporter we observed a significant accumulation of both *Cbx7*-GFP (Fig 7A and B) and H2AK119ub1 signals (Fig 7C and D) at the chromocenters in the Dnmt1i and in the combined OHT and Dnmt1i treated cells compared to DMSO and OHT treatments, without detecting a reduction in corresponding protein levels (Supp fig 5D). Importantly, double treatment with OHT and Dnmt1i led to a significant increase in *Cbx7*-GFP signal at the chromocenters compared to Dnmt1i only (Fig 7B), suggesting that the presence of KDM2b might be sequestering the PRC2 complex away from the PCH and towards newly hypomethylated CGIs (46, 49, 50). This was not the case for H2AK119ub1 where there was no significant difference between Dnmt1i and combined OHT and Dnmt1i treatment (Fig 7D). Taken together these results suggest that while KDM2b is not necessary for the recruitment of PRC1 or PRC2 to the hypomethylated PCH, it might function to direct the complexes to other target regions in the genome.

**Fig 7.**
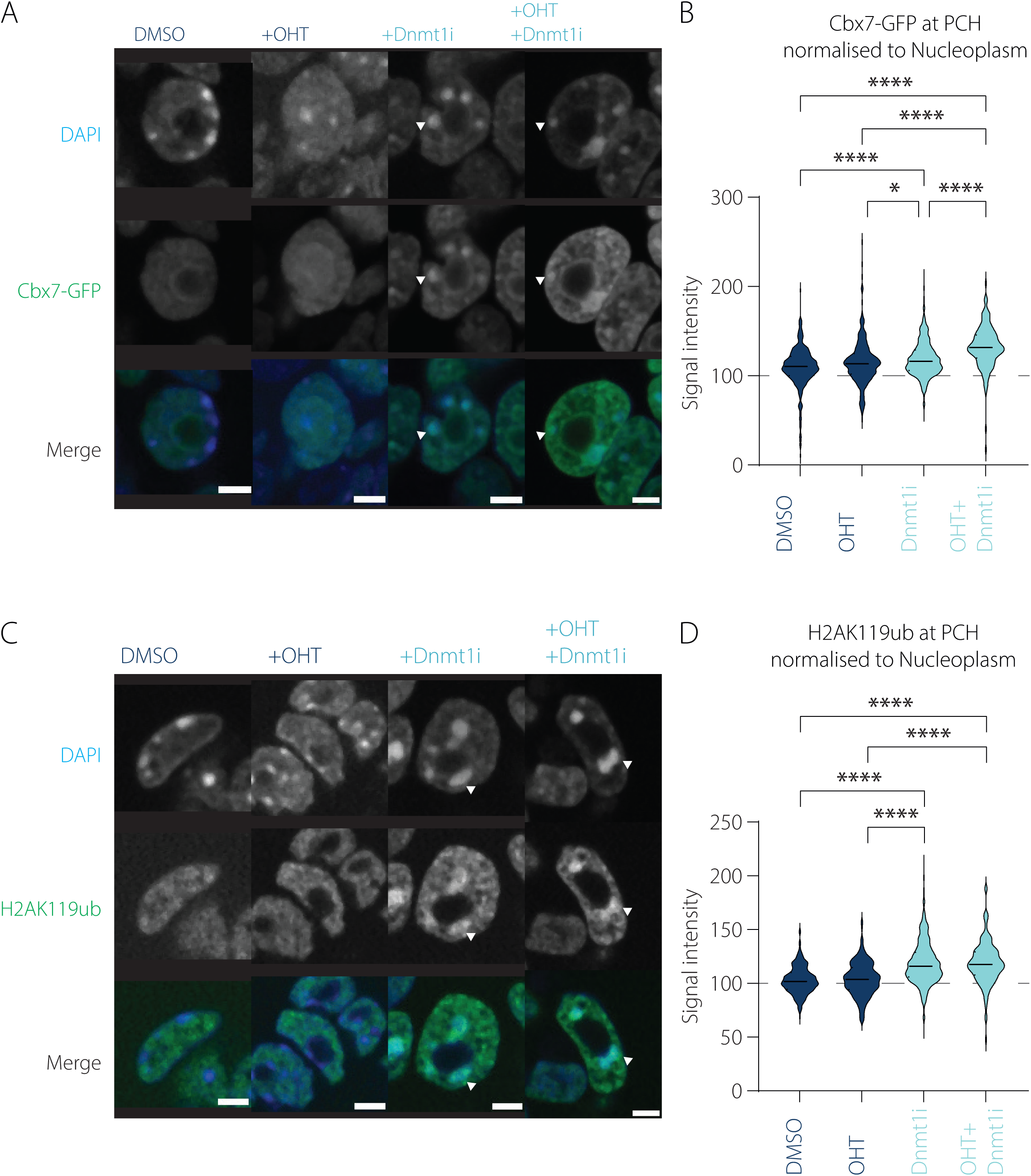
Kdm2b is not necessary for the redistribution of PRC1 or PRC2 to PCH. (A) IF for Cbx7-GFP counterstained with DAPI in Kdm2b cells treated with DMSO, OHT, Dnmt1i or both OHT and Dnmt1i. Representative nuclei from single slice of 3D stack are shown. Z-stacks obtained using 40x air objective. Scale bar corresponds to 5µm. Arrowheads point to a representative chromocenter at which the DAPI and reporter signals overlap. (B) Violin plot of mean signal intensity of Cbx7-GFP reporter signal at chromocenter normalised to that in the nucleoplasm in Kdm2b cells treated with DMSO, OHT, Dnmt1i or both OHT and Dnmt1i. Grey dashed line corresponds the threshold above which the signal is considered increased at the chromocenter compared to the nucleoplasm. Significance was determined by Kruskal-Wallis (one-way Anova) test where *p≤0.05; **** p≤0.0001. (C) As in (A) IF for H2AK119ub counterstained with DAPI in Kdm2b cells treated with DMSO, OHT, Dnmt1i or both OHT and Dnmt1i. Arrowheads point to a representative chromocenter at which the DAPI and antibody signals overlap. (D) As in (B) Violin plot of mean signal intensity of H2AK119ub antibody signal at chromocenter normalised to that in the nucleoplasm in Kdm2b cells treated with DMSO, OHT, Dnmt1i or both OHT and Dnmt1i.

### TET enzymes are not required for the redistribution of PRC to PCH

TET1 has been shown to direct RING1b to chromocenters. This process was then thought to recruit PRC2 and subsequently PRC1 and promote chromocenter clustering (51). Since we also observe larger chromocenters in our Dnmt1 inhibited cells (Supp fig 1 D and E) suggestive of clustering, we hypothesised that the CXXC containing TET enzymes might also play a role in PcG recruitment to these hypomethylated regions. To test this possibility, we used a TET1/2/3 triple knockout (TET TKO) cell line with its corresponding WT control (30). The TET TKO mESCs are normally cultured in combination with MEK inhibition and GSK3 inhibition (2i/LIF) as they cannot proliferate properly in serum/LIF conditions, perhaps due to excess 5mC levels (22). Southern blotting shows a reduction in methylation levels at minor satellite sequences in 2i/LIF wild-type mESC but these sequences are relatively hypermethylated in TET TKO mESCs as evidenced by the higher molecular weight profile for the minor satellite digest pattern (Supp fig 6A). Treatment with the Dnmt1i showed a marked decrease in DNA methylation at minor satellite sequences in both the WT and TET TKO cells (Supp fig 6A) without affecting global levels of H3K27me3 or H2AK119ub1 (Supp fig 6B). We observed a significant increase in H2AK119ub1 foci formation with visibly larger chromocenters in both Dnmt1i treated conditions compared to the DMSO (Fig 8A and B). Introduction of the *Cbx7*-GFP reporter revealed a similar significant accumulation of reporter signal at the foci upon Dnmt1i treatment compared to DMSO in both the WT and TET TKO cells (Fig 8C and D). These results indicate that loss of the TET enzymes does not impact the ability of H3K27me3 or H2AK119ub1 to relocalise to the hypomethylated PCH.

**Fig. 8.**
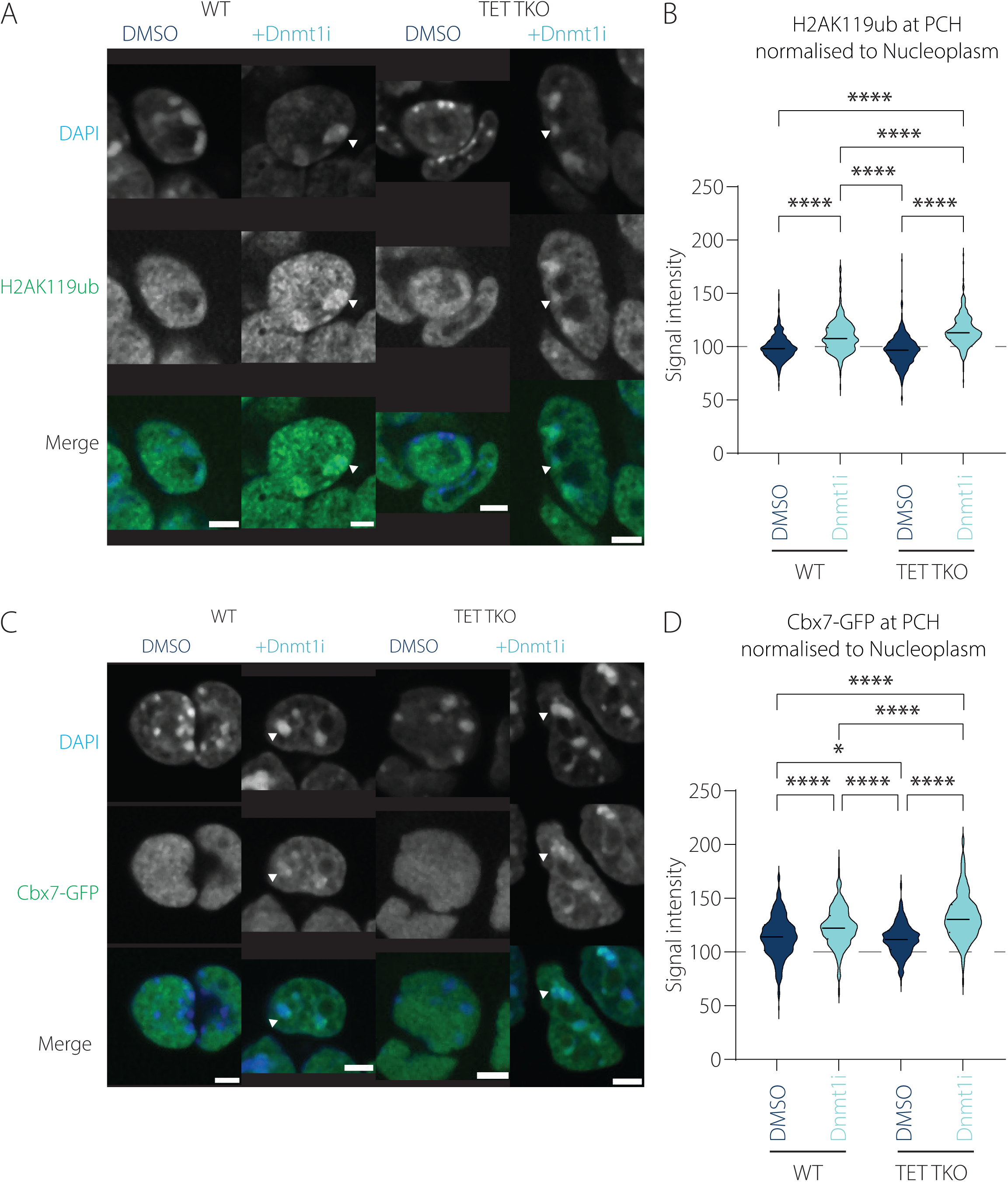
TET enzymes are not required for the redistribution of PRC to PCH. (A) IF for H2AK119ub counterstained with DAPI in TET WT and TKO cells treated with DMSO or Dnmt1i. Representative nuclei from single slice of 3D stack are shown. Z-stacks obtained using 40x air objective. Scale bar corresponds to 5µm. Arrowheads point to a representative chromocenter at which the DAPI and antibody signals overlap. (B) Violin plot of mean signal intensity of H2AK119ub antibody signal at chromocenter normalised to that in the nucleoplasm in TET WT and TKO cells treated with DMSO or Dnmt1i. Grey dashed line corresponds the threshold above which the signal is considered increased at the chromocenter compared to the nucleoplasm. Significance was determined by Kruskal-Wallis (one-way Anova) test where **** p≤0.0001. (C) As in (A) IF for Cbx7-GFP reporter counterstained with DAPI in TET WT and TKO cells treated with DMSO or Dnmt1i. Arrowheads point to a representative chromocenter at which the DAPI and reporter signals overlap. (D) As in (B) Violin plot of mean signal intensity of Cbx7-GFP reporter signal at chromocenter normalised to that in the nucleoplasm in TET WT and TKO cells treated with DMSO or Dnmt1i.

## Discussion

A long-held view is that separate facultative and constitutive heterochromatin organisations provide repressive control of different elements of the epigenome and nuclear architecture of the cell. This view was challenged by findings from the perturbations of histone and DNA methyltransferases, revealing a degree of cooperation between repressive systems in mESCs (52, 53). Our understanding of this interplay and its role is still limited. A phenomenon that has raised much interest is the redistribution of facultative heterochromatin hallmarks, such as Polycomb repressive complexes (PRCs) and the H3K27me3 epigenetic mark, away from developmentally repressed genes and into constitutive heterochromatin regions. This redistribution occurs in mESCs depleted of constitutive heterochromatin components such as H3K9me3 or DNA methylation (54, 55).

### A dynamic, inducible and reversible model system to study PcG relocalisation to PCH

Our knowledge of the heterochromatin redistribution process was gained through genomic mapping technologies, immunofluorescence imaging and molecular approaches. These have been mostly based on comparisons between wild type and established gene knockout cell lines (55, 56). Cellular adaptation to the deletion of essential epigenetic modifiers could potentially mask the native mechanisms in these cells. Studies of the dynamic redistribution and a causal explanation for these events were largely lacking.

In this study, we developed a platform with which to systematically study the dynamics of PcG relocalisation to PCH. We determined the timeframe within which the complexes can be found at the hypomethylated PCH and evaluated the involvement of accessory proteins in promoting PcG targeting. We employed mESCs with an inducible *Dnmt1* knockout or treated with a recently characterised small molecule inhibitor of DNMT1 (40, 57). This approach offers rapid and reversible hypomethylation and therefore avoids secondary effects that might arise from adaptations. Advanced imaging of fluorescently labelled nuclei enabled the observation of the timeline of heterochromatin changes from snapshots of populations of individual nuclei. Using a custom designed image analysis workflow for the data from these nuclei allowed us to quantitate the transitions. We contribute findings observed for the first time in the context of the actual transition between heterochromatin states in mESC by showing that H3K27me3 and H2AK119ub1 relocalisation to PCH is dynamic and coincident with the occurrence of DNA hypomethylation. This localisation at PCH is lost when DNA methylation is reimposed. The similarities and differences from steady state analysis in mESC are informative with regards to the potential mechanisms and relevant to understanding dynamic epigenetic reprogramming in early embryos and induced pluripotent stem cells.

It is important to note that the custom macro, described in this study, is designed to specifically detect the chromocenters that meet the thresholding requirements (described in Methods), which allows for user-independent quantification. Not all chromocenters in the nucleus will meet these requirements as they might be in a different plane to the one that is used for quantification. The large number of nuclei that can be investigated using our macro, with its capability of quantitating variability in the redistribution to PCH, increases the statistical power and sensitivity of detection. Additionally, the macro is capable of detecting signal in the area outside the chromocenters – the nucleoplasm. This is necessary for the definition of the foci, but it also means that while signal might be concentrated in the chromocenters, the modifications remain cytologically unchanged at other genomic loci. This accumulation of modifications at PCH is not associated with a change in their overall levels, as confirmed by western blot analysis.

Our data shows the effective depletion of the DNA methylation signal over 7 days and multiple cell divisions. DNA methylation loss through tet-OFF repression or chemical inhibition is a process in which the copying of 5mC modifications onto newly replicated chromosomes by DNMT1 enzyme is halted, and existing fully or partially methylated chromosomes are dispersed between dividing cells (58). Stochastically, some chromocenters may contain a proportion of methylated DNA during the transition stages, and some transient de novo methylation. Within the population of nuclei, this does not result in the disappearance or change in number of chromocenters, but in many nuclei induces a transient size increase of the foci possibly due to the restructuring of the region. While the presence of H3K9me3 remains detectable in the chromocenters, many nuclei accumulate H3K27me3 signal over all or part of the chromocenters at the same time as the region hypomethylates. Our study confirms the relocalisation of H3K27me3 and H2AK119ub1 to the chromocenters following DNA methylation depletion, in keeping with the findings from wild type and knockout mESC genotypes and despite the underlying differences between these models (57).

### Testing Recruitment of PcG targeting to PCH

Confirming that loss of DNA methylation is sufficient to induce PcG redistribution to the PCH, we then checked for reciprocal recruitment mechanisms of PRC1 and PRC2 complexes at PCH upon DNA demethylation. Current models propose that PRC2 complex recruitment is contingent on recruitment of canonical PRC1 (55, 56). However, in the context of our mESC model, gene knockout/depletion experiments of *Ezh2* and *Ring1b* suggest that the absence of each PRC complex does not prevent the recruitment of the other complex, as evidenced by the presence of epigenetic marks deposited by these complexes (H2AK119ub1 and H3K27me3). We conclude that in hypomethylated heterochromatin, PRC1 and PRC2 complexes can be recruited to DNA independently of each other and despite of their initial absence from this compartment. We therefore tested other proteins with proposed PRC recruitment activities for involvement in the PRC redistribution to DNA hypomethylated heterochromatin.

Our findings suggest that neither SCML2, BEND3, KDM2b nor TET1-3 enzymes are necessary to relocalise Polycomb-dependent chromatin modifications to PCH in hypomethylated cells. It is possible that differences in the genetic background of the mES cells used in different studies contributes to the differences between our study and previous studies (22, 59), or that there is some redundancy between the factors investigated in this study or with other as yet unknown factors, in our system. It is also possible that factors such as BEND3 and KDM2b, which we found to be slightly inhibitory to PcG accumulation at PCH, and the slight enhancement by SCML2 and EZH2, might promote the creation of a Polycomb environment after the initial establishment or help establish such domains at other hypomethylated regions in the genome (43, 44, 46).

While neither PcG complex is known to have a sequence specificity binding motifs, it is agreed that both PRC1 and PRC2 complexes have an affinity for hypomethylated DNA (23, 49, 50). CGIs, for example, which are generally hypomethylated, have been proposed to function in a similar manner to Polycomb Repressive Elements in *Drosophila melanogaster* and have been shown to attract PcGs (57). Global hypomethylation which occurs during cancer or during early embryo and germ cell development are events during which PcG redistribution can be observed (50). In the highly dynamic hypomethylated early embryo heterochromatin environment, the presence of H3K27me3 at PCH, in addition to H3K9me3, is a defining feature and may play an important role in structuring PCH and preserving genomic integrity (60). As we reported previously, H3K27me3 loss at PCH occurs concomitantly to the spatial segregation of the epiblast from primitive endoderm (21, 60).

Recent research has revealed a wide range of PRC 1 and 2 complexes with different components and substoichiometric interactors in different cell types with evidence of independent functions and localisations (55). Together, these findings suggest that our observations of independent PRC1 and PRC2 recruitment in chromocenters of Dnmt1-deficient mESC may be specific to the hypomethylated genomic chromatin environment at the PCH. Our observations support a model in which stem cells operate a number of PRC targeting mechanisms tailored to their dynamic chromatin environments; in the presence of DNA methylation, PcGs require accessory proteins to assist with recruitment to their target sites; at hypomethylated PCH this is compatible with a simple multi-site binding mass action model for the spreading of PRC1 and PRC2 complexes at ‘unmasked’ multiple low affinity regions (61, 62). For example, taken together, the hypomethylated major and minor satellite repeats represent over a million binding sites that are present in long tandem arrays of condensed chromatin at mouse acrocentric chromosomes (63). Essentially this can act as an ‘attractor’ for PRC1/2 activities that can be clearly visualised with antibody and Cbx7-GFP reporter based methods due to the megabase scale of the satellite regions. In this context, this may not be much different from the spreading of H3K27me3 into new regions that has been noted in a number of studies (23, 49, 50).

Based on dynamic non-promoter locations in embryos and stem cells, it was hypothesised that Polycomb complexes constantly scan the whole genome for H3K27me3 deposition to compensate for the transcription repression function of DNA methylation and H3K9me3 (19, 64). PRCs have general DNA binding activity as evidenced by H3K27me3 modifications spread widely across genomic regions, suggesting that H3K27me3 deposition should be seen in terms of PRC residency time rather than recruitment (56).

This study shows that DNA methylation loss prompts PRC1 and PRC2 to relocate autonomously to PCH in mESCs. While no tested factor is essential for their recruitment, these factors accumulate in the novel PCH structure, influencing their association with hypomethylated heterochromatin. Future work can use this live cell readout platform to identify if additional factors are involved in the dynamics of histone modification deposition at the hypomethylated pericentromeric heterochromatin and their relative contributions to its structural maintenance in different cellular contexts.

## Supporting information

Table of UpregulatedgenesBend3

## Acknowledgements

We are grateful for the kind provision of the modified *Dnmt1*^tet/tet^ ES cell line by Richard Chaillet, to the Klose lab for the provision of the *Ring1b*-AID and *Kdm2b*^fl/fl^ cell lines and to Tuncay Baubec for kindly sharing the *Cbx7*-GFP reporter with us. This work would not be possible without the support of the staff and equipment at the Advanced Imaging Resource facility at the IGC, the FACS facility and staff at the IGC and sequencing assistance from the Edinburgh Clinical Research Facility Genetics core.

## Supplementary figure legends

**Supp Fig 1.**
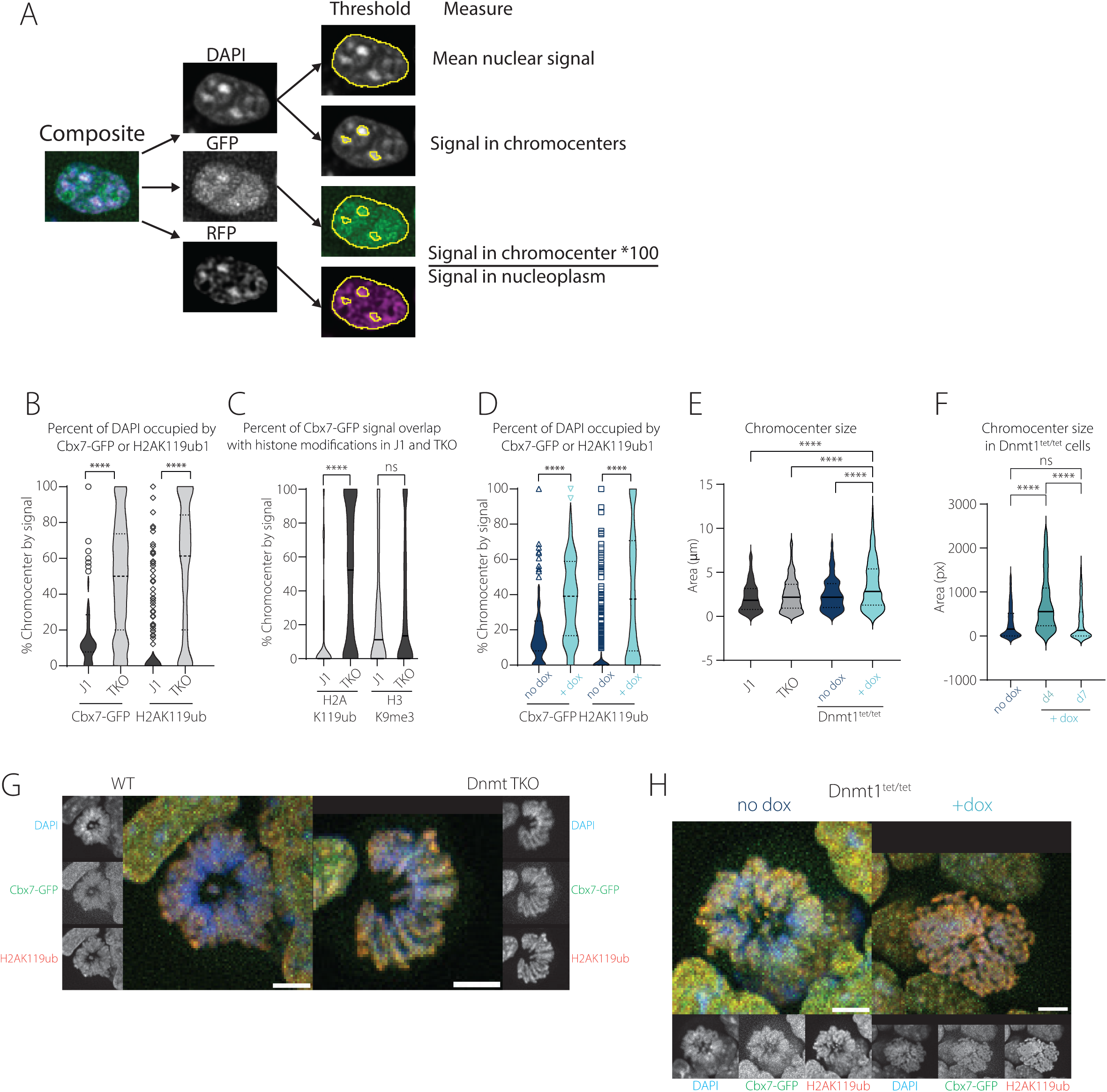
(A) Graphical representation of the macro used for unbiased quantification of signal within the chromocenters as explained in the methods. Briefly, the composite image is split into the individual channels. The DAPI channels is thresholded (Default method) to identify the area of the nucleus. The mean signal intensity is then measured in the DAPI channel and the bottom threshold for the chromocenters is calculated as 1.2 st.dev over the mean signal. The regions of the chromocenters are then subtracted from that of the nucleus to give the nucleoplasm. All regions are then added to ROI manager. The signal intensity in all objects in ROI manager are measured in the rest of the channels. The Mean signal intensity is then calculated as the Mean signal intensity in chromocenter divided by that in the nucleoplasm and multiplied by 100. (B) Violin plot showing the percentage of the chromocenter occupied by increased signal of the reporter/antibody in J1 and TKO cells. Significance was determined by Kruskal-Wallis (one-way Anova) test where **** p≤0.0001 (C) Violin plot of percentage of the Cbx7-GFP defined foci occupied by increased signal of the antibody (H2AK119ub and H3K9me3) in J1 and TKO cells. Significance was determined by Kruskal-Wallis (one-way Anova) test with FDR correction-Benjamini and Hochberg. Test was performed between J1 and TKO samples for each antibody. (D) Violin plot showing the percentage of the chromocenter occupied by increased signal of the reporter/antibody in Dnmt1^tet/tet^ cells with or without dox. Significance was determined by Kruskal-Wallis (one-way Anova) test where **** p≤0.0001 (E) Violin plot showing the chromocenter size (µm) in J1, TKO and Dnmt1^tet/tet^ cells with (4 days) or without dox. Significance was determined by ordinary one-way Anova test with FDR correction-Benjamini and Hochberg. Test was performed between every sample. Non-significant values are not shown. (F) Violin plot of chromocenter size (px) in Dnmt1^tet/tet^ cells in presence of dox for 4 days and 7 days. Significance was determined by ordinary one-way Anova test with FDR correction-Benjamini and Hochberg. Test was performed between every sample. (G) IF on WT and Dnmt TKO cells containing Cbx7-GFP reporter stained for H2AK119ub counterstained with DAPI. Representative mitotic nucleus from maximum intensity Z-projections from 3D stack are shown. C indicates position of centromere. Z-stacks obtained using 40x air objective. Scale bar corresponds to 5µm. (H) IF on Dnmt1^tet/tet^ cells containing Cbx7-GFP reporter stained for H2AK119ub counterstained with DAPI. Representative mitotic nuclei from maximum intensity Z-projections from 3D stack are shown C indicates position of centromere. Z-stacks obtained using 40x air objective. Scale bar corresponds to 5µm.

**Supp fig 2.**
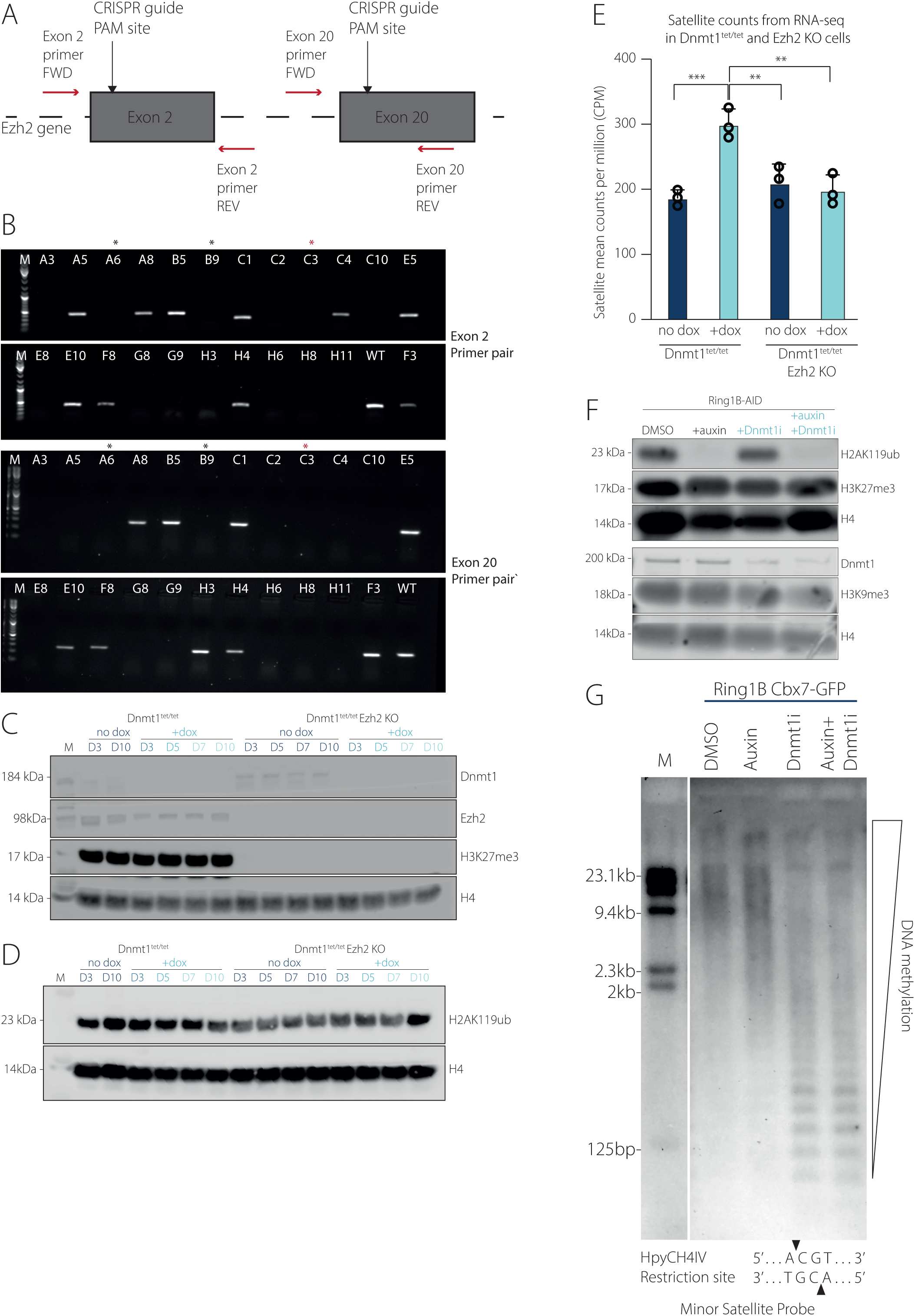
(A) Schematic representation of the CRISPR strategy and sgRNA targeting to generate Ezh2 KO. Adapted from (Lavarone, Barbieri and Pasini, 2019). Primer pairs used for genotyping are shown. CRISPR guide PAM sites are shown to be within the exons. (B) Homozygosity PCR screen using two primer pairs in which one of the primers binds inside the deleted region. Successful Ezh2 KO clones are expected to produce no band while wild type alleles would yield 440bp band Exon 2 primer pair (top two panels) and 290bp band for Exon 20 primer pair (bottom two panels). Selected clone (C3) used in this study is marked with red asterisk * (C) Western blot for Dnmt1, Ezh2, H3K27me3 and H4 (loading control) in Dnmt1^tet/tet^ cells and Ezh2 KO cells with or without dox at different time points. (D) Western blot for H2AK119ub and H4 (loading control) in Dnmt1^tet/tet^ cells and Ezh2 KO cells with or without dox at different time points. (E) Mean satellite read counts per million (CPM) from RNA sequencing in Dnmt1^tet/tet^ cells and Ezh2 KO cells with or without dox. Significance was determined by Ordinary one-way Anova with Tukey’s multiple comparisons test, ** p<0.01, *** p<0.001, **** p<0.0001. (F) Western blot for H2AK119ub, H3K27me3, Dnmt1, H3K9me3 and H4 (loading control) in Ring1b-AID cells treated with DMSO, auxin, Dnmt1i or both auxin and Dnmt1i. (G) Southern blot using methylation sensitive restriction digest (HpyCH4IV enzyme –cut site shown below) followed by hybridisation using probe for minor satellite repeats in Ring1b-AID cells treated with DMSO, auxin, Dnmt1i or both auxin and Dnmt1i.

**Supp fig 3.**
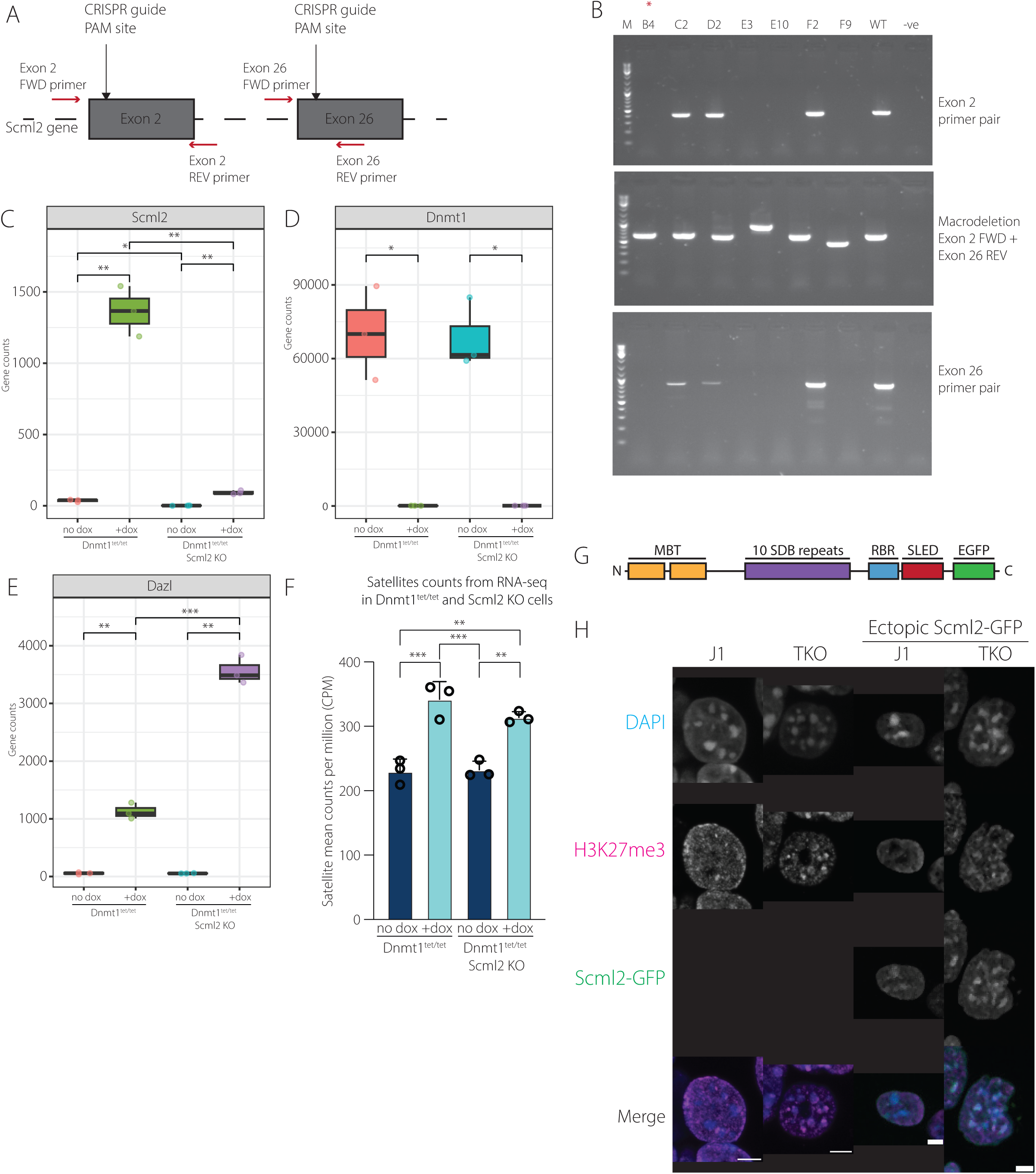
(A) Schematic representation of the CRISPR strategy and sgRNA targeting to generate Scml2 KO. Primer pairs used for genotyping are shown. CRISPR guide PAM sites are shown. (B) PCR screen for homozygous macrodeletion using two primer pairs in which one of the primers binds inside the deleted region. Successful deletion results in no band with the Exon 2 (top panel) and exon 26 (bottom panel) primer pairs. Selected clone (B4) is indicated by red asterisk *. (C) Plot showing normalised read counts of Scml2 gene in Dnmt1^tet/tet^ cells and Scml2 KO cells with or without dox. (D) Plot showing normalised read counts of Dnmt1 gene in Dnmt1^tet/tet^ cells and Scml2 KO cells with or without dox. (E) Plot showing normalised read counts of Dazl gene in Dnmt1^tet/tet^ cells and Scml2 KO cells with or without dox. (F) Mean satellite read counts per million (CPM) from RNA sequencing in Dnmt1^tet/tet^ cells and Scml2 KO cells with or without dox. Significance was determined by Ordinary one-way Anova with Tukey’s multiple comparisons test, ** p<0.01, *** p<0.001. (G) Graphical representation of the Scml2-GFP reporter containing the full Scml2 gene with two Malignant Brain Tumour (MBT) domains, 10 Scml2 DNA binding repeats (SBT), RNA binding region (RBR) domain and Scm-like embedded domain (SLED). EGFP domain was inserted at the C-terminus of the gene. (H) IF for H3K27me3 counterstained with DAPI in parental J1 and TKO as well as in the presence of ectopic Scml2-GFP. Representative nuclei from a max intensity Z-projection of 3D stack are shown. Z-stacks obtained using 60x air objective. Scale bar corresponds to 5µm.

**Supp fig 4.**
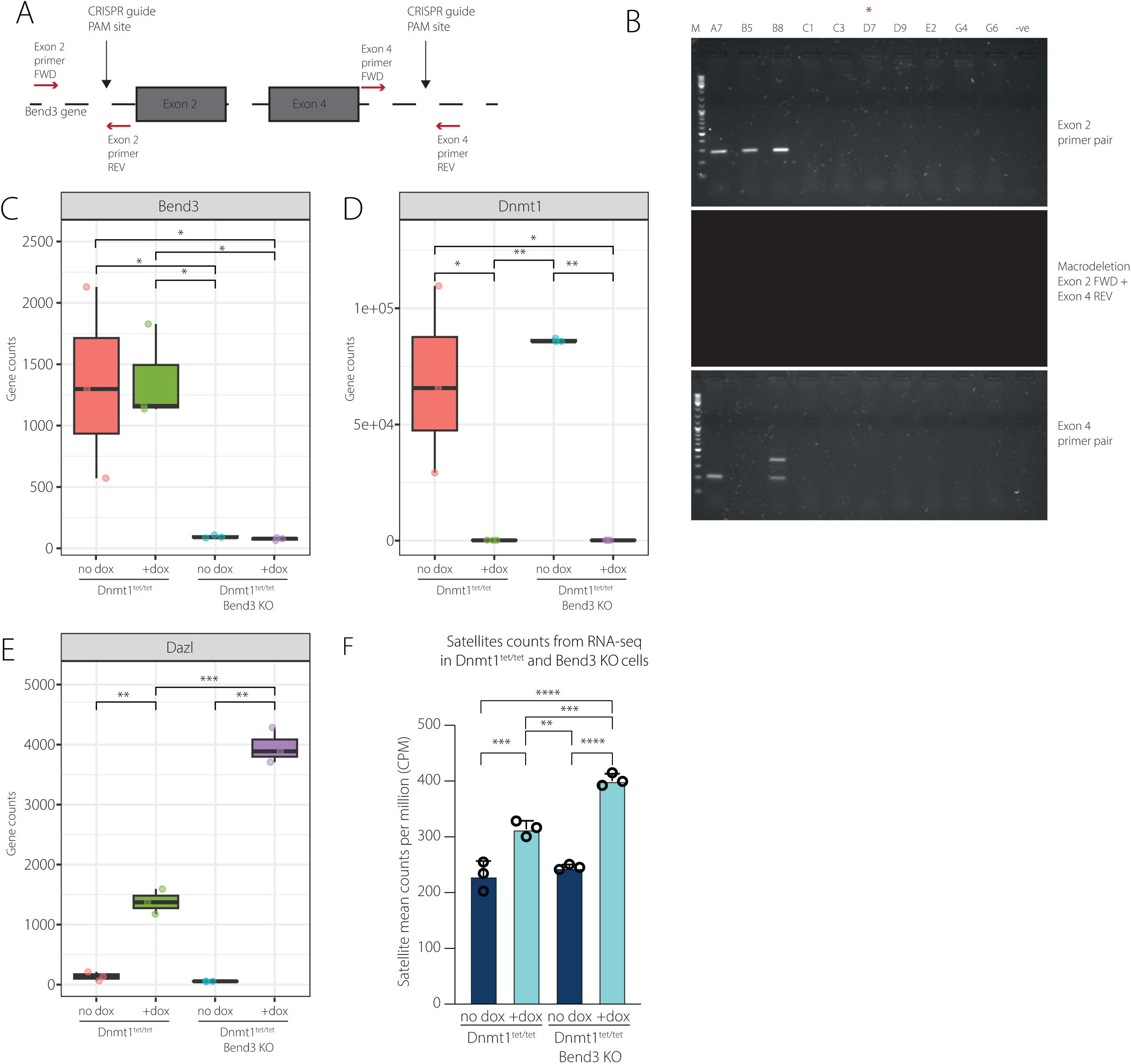
(A) Schematic representation of the CRISPR strategy and sgRNA targeting to generate Bend3 KO. Primer pairs used for genotyping are shown. CRISPR guide PAM sites are shown. (B) PCR screen for homozygous macrodeletion using two primer pairs in which one of the primers binds inside the deleted region. Successful deletion results in no band with the Exon 2 (top panel) and exon 4 (bottom panel) primer pairs. Selected clone (D7) is indicated by red asterisk *. (C) Plot showing normalised read counts of Bend3 gene in Dnmt1^tet/tet^ cells and Bend3 KO cells with or without dox. (D) Plot showing normalised read counts of Dnmt1 gene in Dnmt1^tet/tet^ cells and Bend3 KO cells with or without dox. (E) Plot showing normalised read counts of Dazl gene in Dnmt1^tet/tet^ cells and Bend3 KO cells with or without dox. (F) Mean satellite read counts per million (CPM) from RNA sequencing in Dnmt1^tet/tet^ cells and Bend3 KO cells with or without dox. Significance was determined by Ordinary one-way Anova with Tukey’s multiple comparisons test, ** p<0.01, *** p<0.001, **** p<0.0001.

**Supp fig 5.**
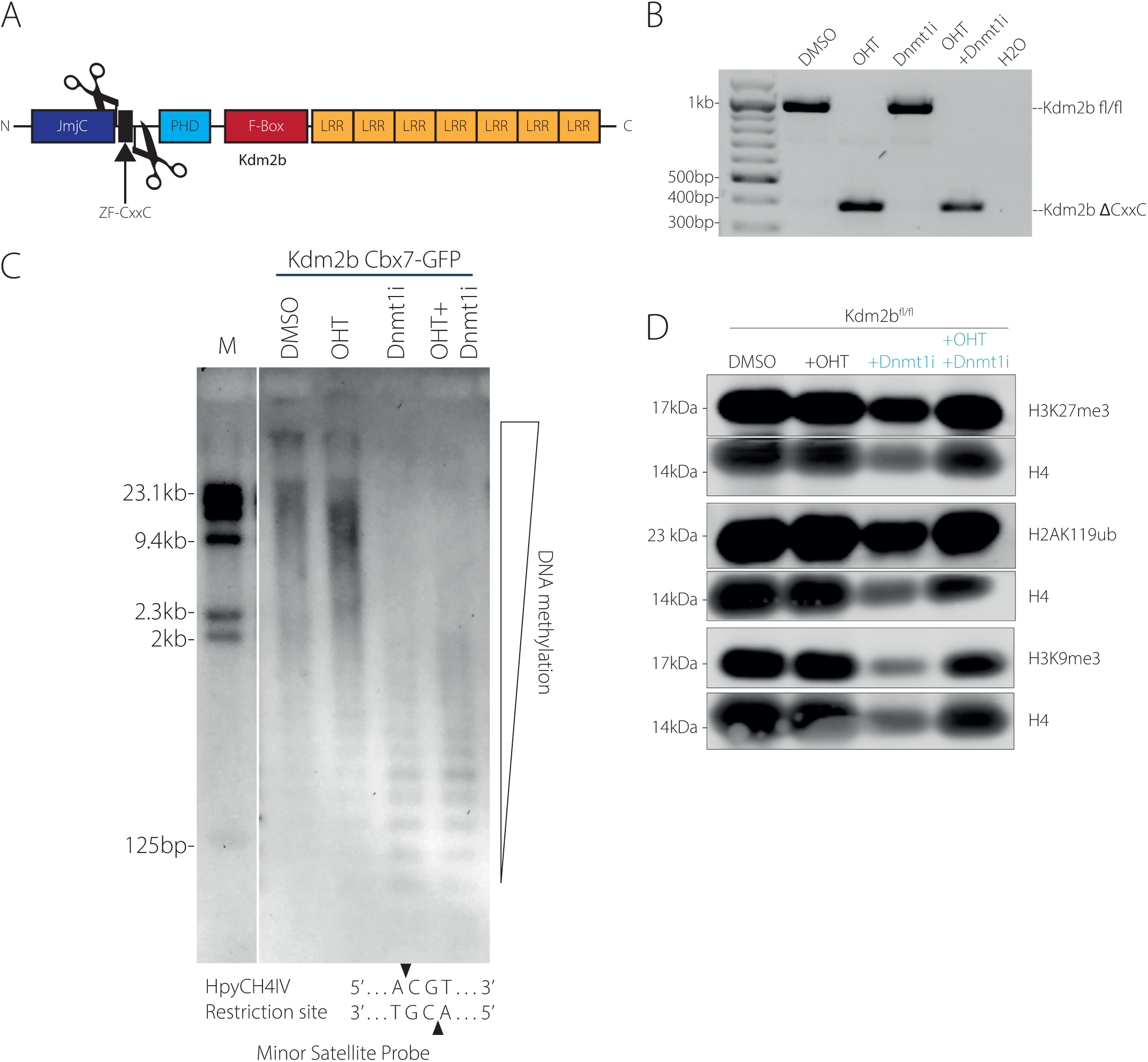
(A) Graphical representation of the Kdm2b protein containing N-terminal JmjC domain, a ZF-CxxC domain, PHD domain, an F-box domain and 7 Leucine-rich repeats (LRR) donated by Klose lab. Graph shows location of LoxP sites (scissors) flanking the ZF-CxxC domain. Upon OHT treatment cre-recombination leads to excision of the domain. (B) PCR with primers flanking the CxxC domain performed on genomic DNA from Kdm2b cells treated with DMSO, OHT, Dnmt1i or both OHT and Dnmt1i. H20 serves as no DNA control. Primers amplify a 1kb fragment in DMSO and Dnmt1i conditions and a smaller (400bp) fragment following recombination induced by OHT. (C) Southern blot using methylation sensitive restriction digest (HpyCH4IV enzyme –cut site shown below) followed by hybridisation using probe for minor satellite repeats in Kdm2b cells containing the Cbx7-GFP reporter treated with DMSO, OHT, Dnmt1i or both OHT and Dnmt1i. Western blot for H2AK119ub, H3K27me3, H3K9me3 and H4 (loading control) in Ring1b-AID cells treated with DMSO, OHT, Dnmt1i or both OHT and Dnmt1i.

**Supp Fig. 6.**
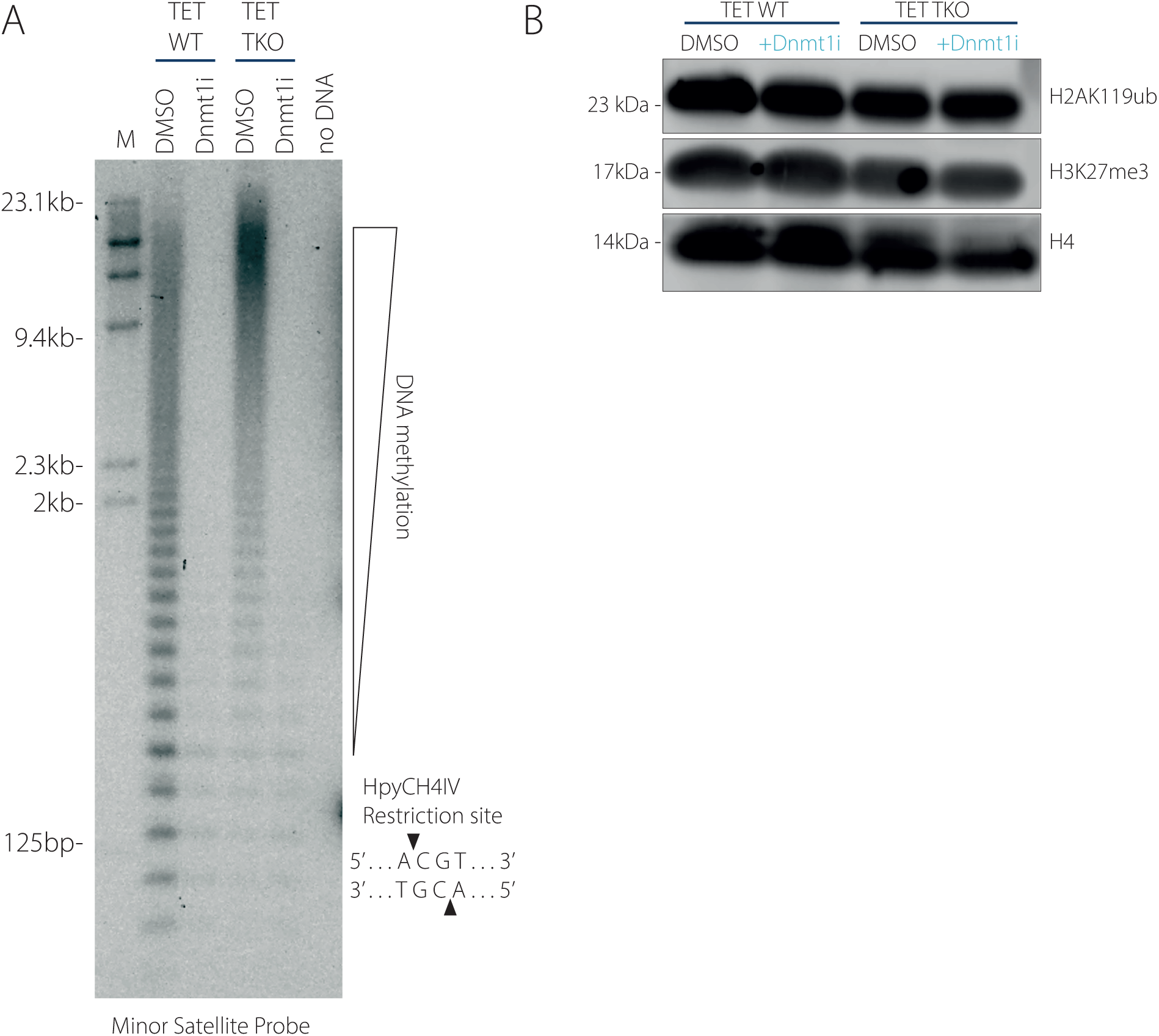
(A) Southern blot using methylation sensitive restriction digest (HpyCH4IV enzyme –cut site shown below) followed by hybridisation using probe for minor satellite repeats in TET WT and TKO cells treated with DMSO or Dnmt1i. (B) Western blot for H2AK119ub, H3K27me3 and H4 (loading control) in TET WT and TKO cells treated with DMSO or Dnmt1i.

